# Lupus IgA1 autoantibodies synergize with IgG to enhance pDC responses to RNA-containing immune complexes

**DOI:** 10.1101/2023.09.07.556743

**Authors:** Hayley R. Waterman, Matthew J. Dufort, Sylvia E. Posso, Minjian Ni, Lucy Z. Li, Chengsong Zhu, Prithvi Raj, Kelly D. Smith, Jane H. Buckner, Jessica A. Hamerman

## Abstract

Autoantibodies to nuclear antigens are hallmarks of the autoimmune disease systemic lupus erythematosus (SLE) where they contribute to pathogenesis. However, there remains a gap in our knowledge regarding how different isotypes of autoantibodies contribute to disease, including the production of the critical type I interferon (IFN) cytokines by plasmacytoid dendritic cells (pDCs) in response to immune complexes (ICs). We focused on IgA, which is the second most prevalent isotype in serum, and along with IgG is deposited in glomeruli in lupus nephritis. Here, we show that individuals with SLE have IgA autoantibodies against most nuclear antigens, correlating with IgG against the same antigen. We investigated whether IgA autoantibodies against a major SLE autoantigen, Smith ribonucleoproteins (Sm/RNPs), play a role in IC activation of pDCs. We found that pDCs express the IgA-specific Fc receptor, FcαR, and there was a striking ability of IgA1 autoantibodies to synergize with IgG in RNA-containing ICs to generate robust pDC IFNα responses. pDC responses to these ICs required both FcαR and FcγRIIa, showing a potent synergy between these Fc receptors. Sm/RNP IC binding to and internalization by pDCs were greater when ICs contained both IgA1 and IgG. pDCs from individuals with SLE had higher binding of IgA1-containing ICs and higher expression of FcαR than pDCs from healthy control individuals. Whereas pDC FcαR expression correlated with blood ISG signature in SLE, TLR7 agonists, but not IFNα, upregulated pDC FcαR expression in vitro. Together, we show a new mechanism by which IgA1 autoantibodies contribute to SLE pathogenesis.

**One Sentence Summary:** IgA1 autoantibodies synergize with IgG in RNA-containing immune complexes to generate robust pDC IFNα responses in a FcαR receptor dependent manner.

## INTRODUCTION

Systemic lupus erythematosus (SLE) is an autoimmune disease characterized by the breakdown of tolerance to nuclear antigens and systemic inflammation. Individuals with SLE have circulating autoreactive anti-nuclear antibodies (ANAs) that recognize nuclear antigens such as dsDNA and DNA or RNA-associated proteins, such as histones or subunits of ribonucleoproteins (RNPs) (*1–3*). ANAs form nucleic acid-containing immune complexes (ICs), which deposit in tissues where they are recognized by and trigger inflammatory responses in Fc receptor (FcR) expressing cells such as dendritic cells, monocytes, and neutrophils (*4–8*). In this way ANA-ICs facilitate nucleic acid entry to endosomes where RNA and DNA sensing innate immune receptors reside, thereby allowing immune cells to aberrantly respond to autologous RNA and DNA molecules.

While it is generally accepted that autoantibodies to nuclear antigens contribute to SLE pathology, there is still much to be discovered regarding how different isotypes and antigen specificities contribute to disease. Most studies have focused on IgG isotype ANAs in SLE (*1, 9*), however there are four additional isotypes present in serum: IgA, IgM, IgD and IgE (*10–12*). IgA is the second most prevalent antibody isotype after IgG (*10*), and there is some evidence that IgA ANAs may contribute to SLE pathology. A few studies have shown that IgA anti-dsDNA antibodies are present in 30-50% of individuals with SLE and that IgA anti-dsDNA antibodies correlated to SLE disease activity index (SLEDAI), lupus nephritis, and other markers of SLE severity, such as high erythrocyte sedimentation rate and low complement C3 and C4 (*13–15*). IgA is almost uniformly present in lupus nephritis glomeruli, in addition to IgG, IgM, C3 and C1q, and is used for diagnosis (*16*). Elevated levels of IgA ANAs have also been found in the saliva and fecal samples of SLE subjects (*17, 18*). Additionally, one study found a substantial increase in IgA isotype B cell clones in SLE compared to healthy controls and other immune-mediated diseases (*19*). However, the pathophysiologic significance of IgA in lupus nephritis is not understood and requires further investigation. Several studies have shown that IgA antibodies have a pathogenic role in other inflammatory diseases e.g., IgA nephropathy, IgA vasculitis, dermatitis herpetiformis, inflammatory bowel disease, linear IgA bullous disease and rheumatoid arthritis (*20*).

IgA is a unique isotype that has multiple forms and functions. IgA is generally recognized as important in mucosal immunity where it is secreted as a secretory dimeric (SIgA) form that can bind and neutralize pathogens (*20*). IgA is also abundant in human serum, where it exists mainly in a monomeric form that can be regulatory and promote immune homeostasis (*20–22*). The IgA-specific FcαR (CD89) associates with the immune tyrosine activating motif (ITAM)-containing FcR common gamma (FcRγ) chain to confer downstream signaling and internalization in response to IgA-FcαR binding (*23, 24*). When monomeric IgA binds to FcαR, the associated ITAMs are only partially phosphorylated resulting in recruitment of SHP-1 and downstream inhibitory signaling (*25–27*). Multimeric or complexed IgA binding to FcαR causes clustering of the receptor and the associated FcRγ chains resulting in complete phosphorylation of FcRγ ITAMs, recruitment of Syk, and downstream pro-inflammatory signaling (*24, 28*). In this way, IgA functions as inhibitory when monomeric and stimulatory when multimeric. In some contexts, IgA-FcαR interactions are more effective than IgG-FcγR interactions at stimulating immune responses. Multimeric IgA can be more potent than IgG at activating neutrophils (*29, 30*) and FcαR signaling enhances toll-like receptor (TLR) signaling in monocytes and macrophages (*31*). Lastly, a non-synonymous SNP in *FCAR*, encoding FcαR, which enhances signaling through this receptor, was associated with increased risk of SLE in a case-control study (*32*). Thus, there is evidence that IgA has diverse roles in immune responses, and may be important in SLE. Because there is no mouse orthologue of *FCAR*, studies of IgA and FcαR in human disease require human systems.

Because of the emerging studies suggesting IgA autoantibodies may be relevant in SLE and data demonstrating that multimeric IgA is potent at stimulating immune cells through FcαR, we wanted to better understand the role IgA ANAs play in the context of SLE. Specifically, we asked how IgA ANAs contribute to plasmacytoid dendritic cell (pDC) responses in SLE. pDCs are specialized innate immune cells implicated in SLE pathogenesis that release large amounts of the key type I interferon (IFN) cytokines in response to RNA and DNA sensing by TLRs 7 and 9, respectively (*8, 33–36*). Type I IFNs are cytokines critical for anti-viral immune responses with myriad immune modulating functions, including promoting B cell survival, plasma cell differentiation, T cell proliferation and DC maturation (*37*). Additionally, type I IFNs promote the expression of interferon-stimulated genes (ISGs) in many cell types, which are upregulated in individuals with SLE compared to healthy control individuals and correlate to measures of SLE disease severity (*3, 38, 39*). In pDCs, FcR recognition of ANA-ICs facilitates the internalization of nucleic acids to endosomes, where TLR7 and TLR9 reside (*5, 8*). This leads to TLR signaling via IRF7 and subsequently potent type I IFN production, contributing to the pathogenesis of SLE (*39, 40*).

Given the important role of pDCs in type I IFN production in SLE, we chose to focus on the effect IgA-containing ICs have on these cells. We found pDCs express FcαR in addition to the IgG-binding FcγRIIa and can bind both IgA and IgG. Using a unique in vitro system employing serum from individuals with SLE to generate ICs with potent pDC stimulatory function, we show here a novel and critical function for the IgA1 subtype in pDC type I IFN production in response to RNA-containing RNPs. FcαR and FcγRIIa acted synergistically to induce type I IFN production through facilitating internalization of ICs, and pDCs from individuals with SLE had higher IC internalization and FcαR expression than those from HCs. Thus, we have uncovered a critical role of IgA1 and FcαR in pDC responses in SLE.

## RESULTS

### SLE donors have IgA isotype ANAs to multiple nuclear autoantigens

To better understand the breadth and prevalence of these IgA anti-nuclear antibodies (ANA), we performed autoantigen profiling on serum from 24 SLE subjects assessing both IgA and IgG isotypes. Supplemental Table 1 summarizes the demographics and clinical features including disease activity scores (SLEDAI), presence of autoantibodies (ANA, dsDNA, anti-Smith RNP) and disease duration. We found that all 24 SLE donors had detectable IgA isotype autoantibodies against multiple RNA-associated and DNA-associated nuclear antigens (Fig. 1A, Supplementary Fig. 1A). When comparing IgA and IgG autoantibodies to the same autoantigen, we found that the level of these antibodies positively correlated in 25/26 nuclear antigens relevant to SLE, whereas the degree of correlation and the ratio of IgA to IgG autoantibody varied by antigen (Fig. 1A, Supplementary Fig. 1A and 1B). Antibodies against Smith/ribonucleoprotein (Sm/RNP) and double stranded DNA (dsDNA) are prevalent in SLE (*41, 42*) and IgA and IgG autoantibodies against these antigens were two of the most highly correlated of all nuclear antigens tested (Fig. 1B and Supplementary Fig. 1B).

**Figure 1.**
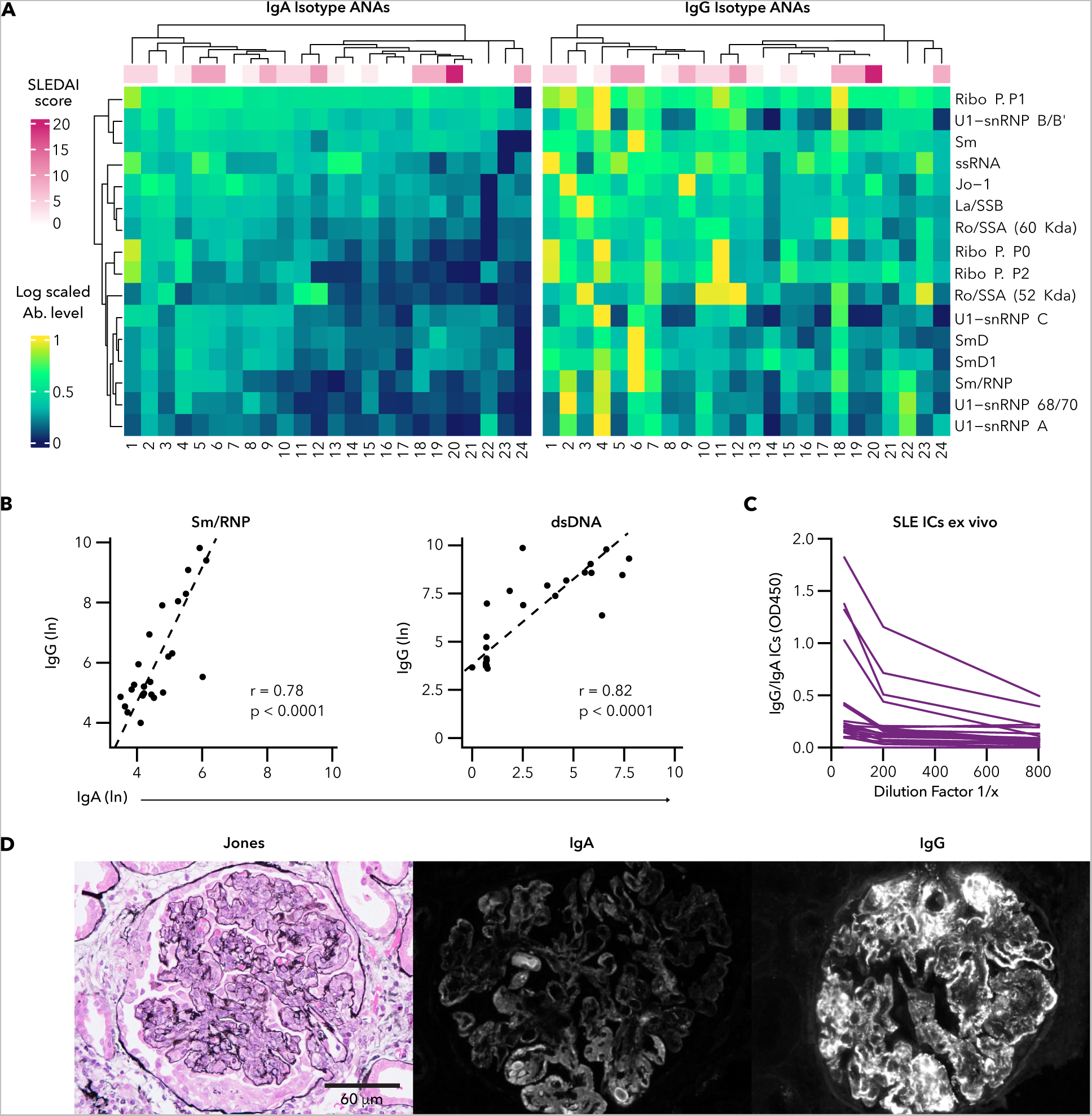
IgA and IgG RNA-associated anti-nuclear antibodies in SLE serum. **(A)** Autoantibody profiling of serum from 24 SLE subjects. Heatmaps of log-scaled antibody binding for IgA (*left*) and IgG (*right*) isotype antibodies to 16 RNA-associated nuclear antigens. Each row corresponds to an antigen and each column corresponds to an individual subject. Color bar on top indicates SLEDAI score for each donor at time of blood draw. Clustering of subjects and samples is based on similarity in scaled IgA levels, with the same clustering applied to the IgG heatmap. (**B**) Correlation between IgA and IgG for anti-Sm/RNP (*left*) and dsDNA (*right*) nuclear antigens. Each symbol represents an individual subject. **(C)** Serial dilution curves of immune complexes (ICs) containing both IgA and IgG in serum from 26 SLE subjects as measured by ELISA. Of the 26 SLE subjects 15% (4/26) had no detectable IgA/IgG ICs. Each line represents an individual subject. **(D)** Histology and immunofluorescence of representative glomeruli from class IV lupus nephritis showing Jones silver staining (*left*), and IgA (*middle*) and IgG (*right*) staining of serial sections. (**B**) r = Pearson correlation coefficient.

As an individual’s serum contained both IgA and IgG autoantibodies to the same nuclear antigens, we reasoned that the immune complexes (ICs) formed in vivo when these autoantibodies recognize their cognate nuclear antigens will likely contain both IgG and IgA isotype antibodies. To test whether circulating ICs with both IgA and IgG are found in SLE, we performed ELISAs on SLE serum (n=26) using anti-IgG capture antibody and anti-IgA detection antibody. In this way we could detect ICs that contained both IgA and IgG regardless of antigen specificity. We found that 21/26 SLE donors we tested had detectable mixed IgA/IgG ICs directly ex vivo (Fig. 1C). As has been well established (*16*), we also detected IgA in the glomeruli of diffuse proliferative (class IV) lupus nephritis (Fig. 1D). The fact that mixed IgA/IgG ICs exist in SLE serum, that SLE serum often contains both IgA and IgG isotype ANAs, and that lupus nephritis usually shows both IgA and IgG deposition, led us to hypothesize that IgA/IgG mixed isotype ICs may contribute to disease pathogenesis in SLE. Because IgA and IgG isotype antibodies can have different effector functions and are recognized by different FcRs, these isotypes have the potential to both contribute to pathogenic IC-mediated responses in SLE.

### pDCs express the IgA-specific FcαR and bind IgA

Because pDCs make large quantities of IFNα in response to ICs, we wanted to determine how IgA-containing ICs contribute to the responses of pDCs. pDCs express the IgG-specific FcγRIIa (CD32a) and respond to IgG-containing ICs, however FcαR expression on or IgA binding to pDCs has not been described. Therefore, we first examined whether pDCs express the IgA-specific FcαR (CD89) and bind to IgA. Analysis of publicly available single-cell RNA sequencing data (*43*) revealed that pDCs expressed *FCAR* mRNA that encodes the IgA-specific FcαR, albeit to a lesser degree than monocytes, which are known to have high FcαR expression (Supplementary Fig. 2A). Interestingly, a higher percentage of pDCs had detectable *FCAR* mRNA than *FCGRIIA* mRNA encoding FcγRIIa, which also had lower percentage and mean counts of expression in pDCs compared to monocytes. We confirmed that pDC expression of *FCAR* mRNA resulted in FcαR protein surface expression by flow cytometry (Fig. 2A, Supplementary Fig. 2B). We also observed FcγRIIa surface expression on pDCs and saw that both FcαR and FcγRIIa surface expression was lower on pDCs compared to monocytes, which was consistent with the scRNA-Seq data (Fig. 2A, 2B, Supplementary Fig. 2B-C). To determine if pDCs could bind to IgA, we heat-aggregated biotinylated human IgA and used fluorescently-labeled streptavidin to visualize the IgA by flow cytometry. In this assay, pDCs bound heat aggregated IgA (Fig. 2C). In addition, we detected bound IgA on the surface of pDCs directly ex vivo by staining pDCs with anti-IgA antibody (Fig. 2D). As expected, pDCs also bound heat-aggregated IgG in vitro and had surface bound IgG directly ex vivo (Fig. 2C, 2D) Therefore, pDCs have the potential to respond to both IgA and IgG through FcαR and FcγRIIa, respectively. As we showed above that individuals with SLE have circulating ICs containing both IgA and IgG, we hypothesized that these receptors act in concert in pDCs to recognize ICs.

**Figure 2.**
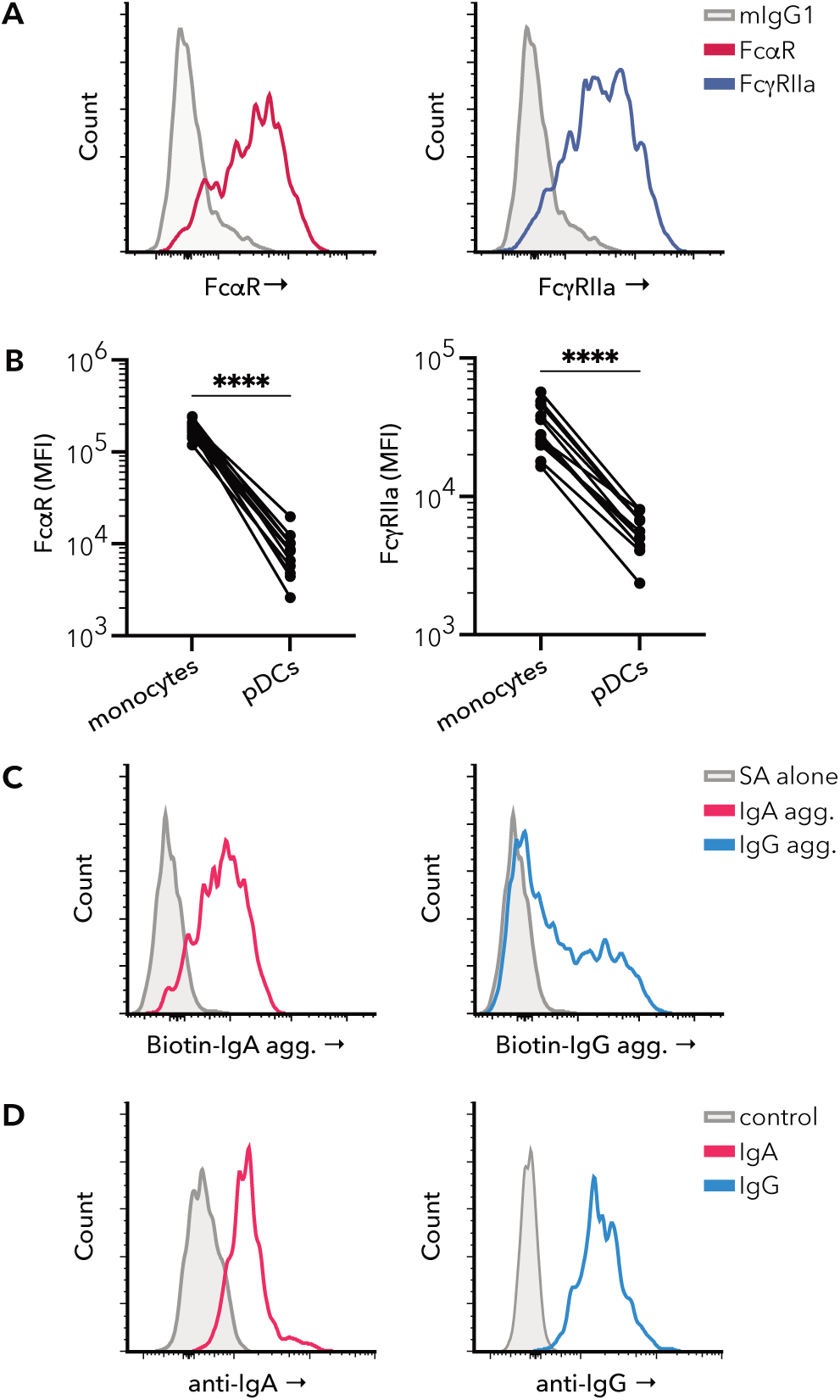
Human blood pDCs express FcαR and FcγRIIa and bind IgA and IgG. Surface staining and flow cytometry analysis performed on PBMCs from healthy control (HC) donors. **(A)** Representative histograms of FcαR (*red*) and FcγRIIa (*blue*) expression on pDCs compared to mIgG1 isotype control (*grey*), with detection by the same secondary antibody. **(B)** Paired analysis between monocyte and pDC FcαR (left) and FcγRIIa (right) surface staining (n=12 HC donors). **(C)** Binding of IgA and IgG to pDCs was assessed using biotinylated human heat-aggregated IgA or heat-aggregated IgG and fluorescently labeled streptavidin for visualization by flow cytometry, Streptavidin (SA) alone (*grey*), heat-aggregated IgA (*red*) and heat-aggregated IgG (*blue*). **(D)** Direct ex vivo binding of IgA and IgG to pDCs was assessed by staining with anti-IgA (*red*) or anti-IgG antibody (*blue*) directly ex vivo. Control (*grey*) has no anti-IgA or anti-IgG detection antibody. (**A-D**) Staining was performed on thawed PBMCs and pDCs gated as shown in Supplementary Fig. 2B. **(B)** Ratio paired t-test (**** p<0.0001).

### IgA1 in Sm/RNP immune complexes enhances pDC IFNα responses via FcαR and FcγRIIa co-engagement

We next investigated whether IgA in ICs influences pDC production of IFNα. We focused on the IgA1 subtype as this represents the majority of serum IgA (*44, 45*). We modified a previously described in vitro system from *Lood et al* (*46*) to test the effect of IgA1 autoantibodies on pDC IFNα production in response to ICs in vitro. Specifically, we used SLE serum to compare ICs that contained both IgG and IgA1 autoantibodies (SLE serum) to those depleted of IgA1 (SLE serum ΔIgA1) and those generated from purified IgA1 (SLE IgA1) (Fig. 3A). To generate these serum reagents, we first identified SLE donors in our biorepository whose serum contained both IgA and IgG specific for the RNA-containing Sm/RNP nuclear antigen. We found that 75% (12/16) of SLE donors with anti-Sm/RNP IgG also had detectable anti-Sm/RNP IgA by ELISA (Table 1, Supplementary Table 2). To generate IgA1-depleted serum (SLE serum ΔIgA1) that retained IgG and other antibody isotypes, we performed affinity chromatography on SLE serum containing anti-Sm/RNP IgA using Jacalin agarose, which binds to IgA1 (Fig. 3A). Depletion of IgA1 from serum resulted in significant depletion of IgA anti-Sm/RNP antibodies but did not alter the amount of anti-Sm/RNP IgG or IgE as measured by ELISA (Fig. 3B, Supplementary Fig. 3A-B). Purified IgA1 was generated by eluting the IgA1 bound to the affinity column used for generating IgA1-depleted serum. Purified IgA1 had no detectable contaminating IgG, whether specific for Sm/RNP or when measuring total IgG (Fig. 3B, Supplementary Fig. 3A-B). IgA1 depletion also did not significantly affect total or anti-Sm/RNP-specific IgE levels, and purified IgA1 contained minimal anti-Sm/RNP-specific IgE (Supplementary Fig. 3A and 3C).

**Figure 3.**
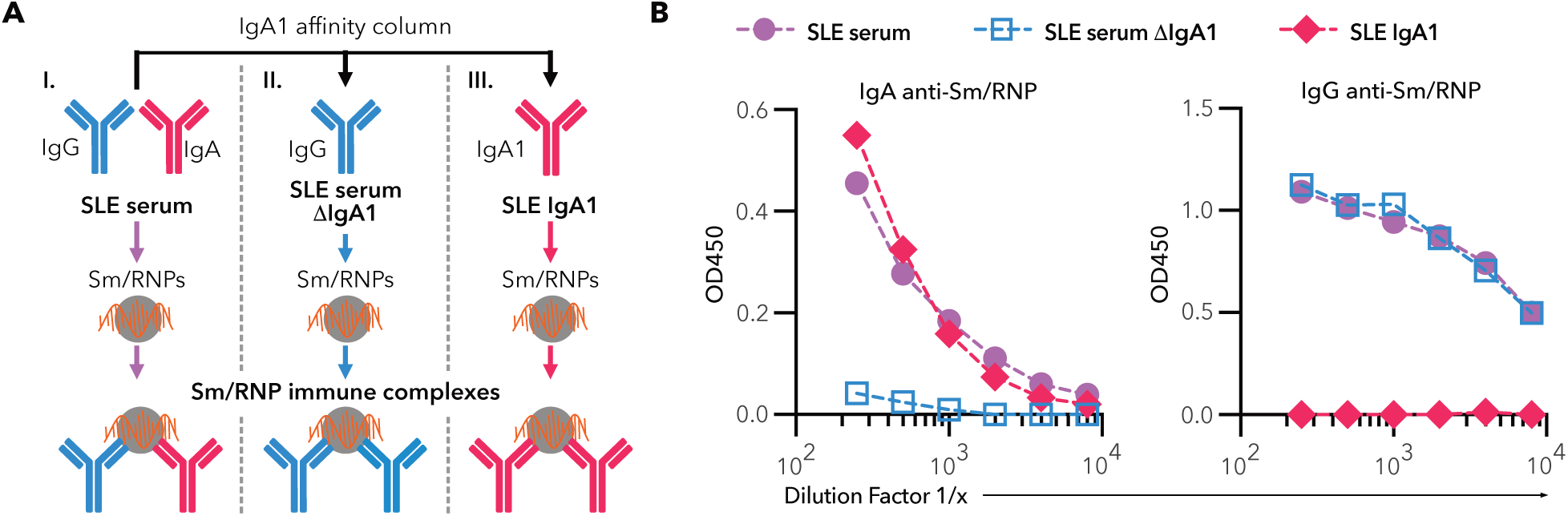
Reagents for generating RNA-containing Sm/RNP immune complexes. **(A)** Schematic showing reagents used to generate immune complexes (ICs) that contained both IgG and IgA (*I.*), mainly IgG (*II.*) or only IgA1 (*III.*) from SLE donors with IgG and IgA anti-Sm/RNP antibodies. IgA1 depleted serum (SLE serum ΔIgA1) and purified SLE IgA1 were generated from SLE serum by using an IgA1 binding substrate column and collecting the flow through and eluted protein, respectively. ICs were made by mixing the different antibody reagents with RNA-containing Sm/RNPs. **(B)** Levels of IgA anti-Sm/RNP (*left*) and IgG anti-Sm/RNP (*right*) antibodies in SLE serum (*purple circles*), SLE serum ΔIgA1 (*blue open squares*) and SLE IgA1 (*red diamonds*), as measured by ELISA. Representative data for one serum donor is shown.

**Table 1.**
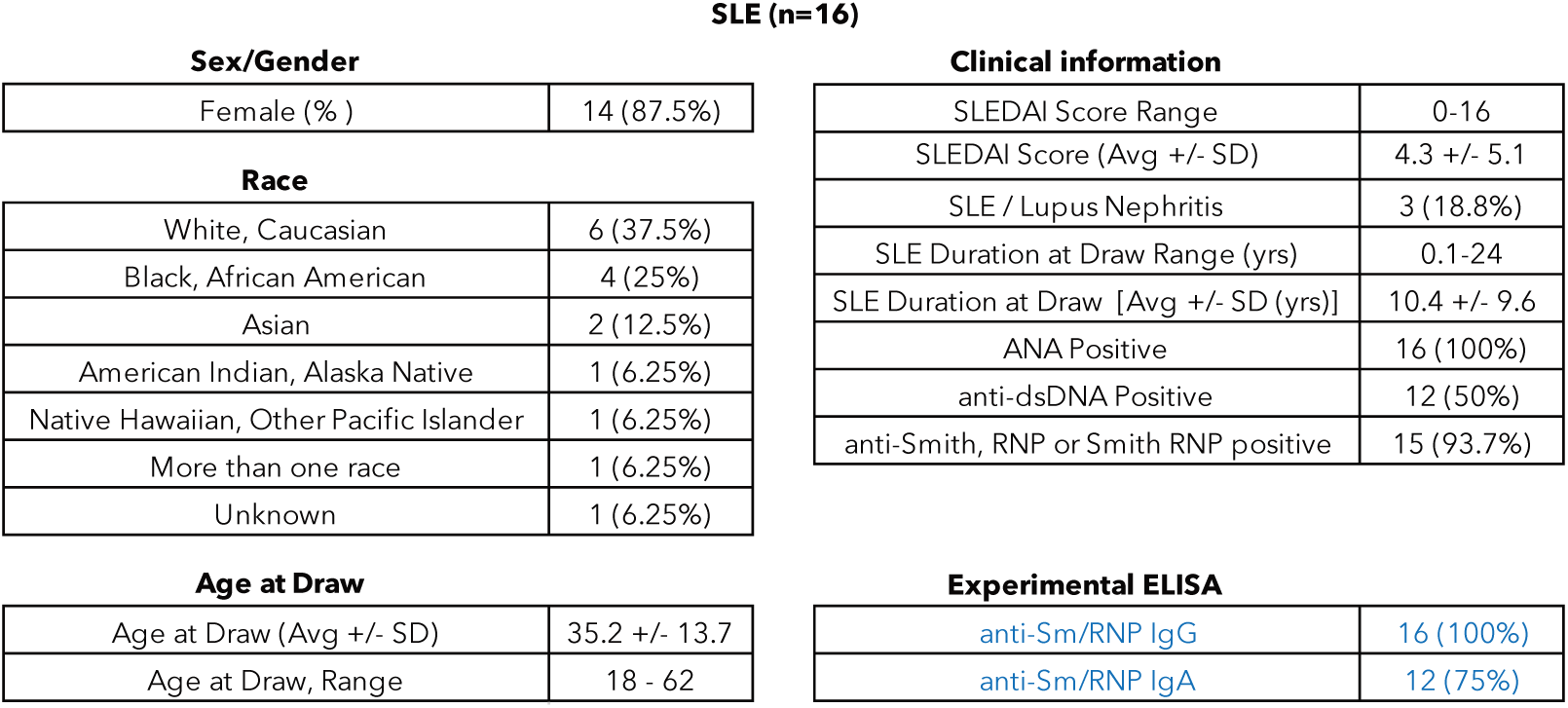
SLE Serum donors.

**SLE (n=16)**
| **Sex/Gender** |  | **Clinical information** |  |
| --- | --- | --- | --- |
| Female (%) | 14 (87.5%) | SLEDAI Score Range | 0-16 |
| **Race** |  | SLEDAI Score (Avg +/- SD) | 4.3 +/- 5.1 |
| White, Caucasian | 6 (37.5%) | SLE / Lupus Nephritis | 3 (18.8%) |
| Black, African American | 4 (25%) | SLE Duration at Draw Range (yrs) | 0.1-24 |
| Asian | 2 (12.5%) | SLE Duration at Draw [Avg +/- SD (yrs)] | 10.4 +/- 9.6 |
| American Indian, Alaska Native | 1 (6.25%) | ANA Positive | 16 (100%) |
| Native Hawaiian, Other Pacific Islander | 1 (6.25%) | anti-dsDNA Positive | 12 (50%) |
| More than one race | 1 (6.25%) | anti-Smith, RNP or Smith RNP positive | 15 (93.7%) |
| Unknown | 1 (6.25%) | **Experimental ELISA** |  |
| **Age at Draw** |  | anti-Sm/RNP IgG | 16 (100%) |
| Age at Draw (Avg +/- SD) | 35.2 +/- 13.7 | anti-Sm/RNP IgA | 12 (75%) |
| Age at Draw, Range | 18 - 62 |  |  |

Using these serum reagents, we generated ICs by combining commercially available Sm/RNPs with either whole SLE serum or IgA1-depleted SLE serum to generate ICs containing both IgG and IgA or principally IgG, respectively (Fig. 3A). These ICs were added to pDCs enriched from fresh healthy control (HC) donor blood, and we measured IFNα secretion in response to the RNA contained within the Sm/RNPs by bioassay. As expected, ICs generated by mixing SLE serum with Sm/RNPs induced a large amount of type I IFNα secretion from HC pDCs from 4 donors, while Sm/RNPs alone elicited very low type I IFNα production (Fig. 4A). Depletion of IgA1 from serum before generation of ICs dramatically abrogated pDC IFNα secretion, which was rescued when the purified IgA1 was added back to IgA1-depleted serum before IC generation (Fig. 4A). Interestingly, ICs generated from purified IgA1 were not sufficient to generate strong pDC IFNα secretion, nor were ICs generated from twice the amount of IgA1-depleted serum, suggesting that IgA1 synergizes with IgG in ICs for pDC responses (Fig. 4A, Supplementary Fig. 4A). To further validate the role of IgA1 in ICs, we purified total IgG and total IgA1 from serum of the SLE donor assessed in Figure 4A, allowing us to generate ICs without additional serum components. Sm/RNP ICs containing purified IgA1 and IgG induced significantly stronger IFNα responses than those induced by ICs made with purified IgG alone (Fig. 4B), consistent with a synergistic response with IgG and IgA1 both present in ICs.

**Figure 4.**
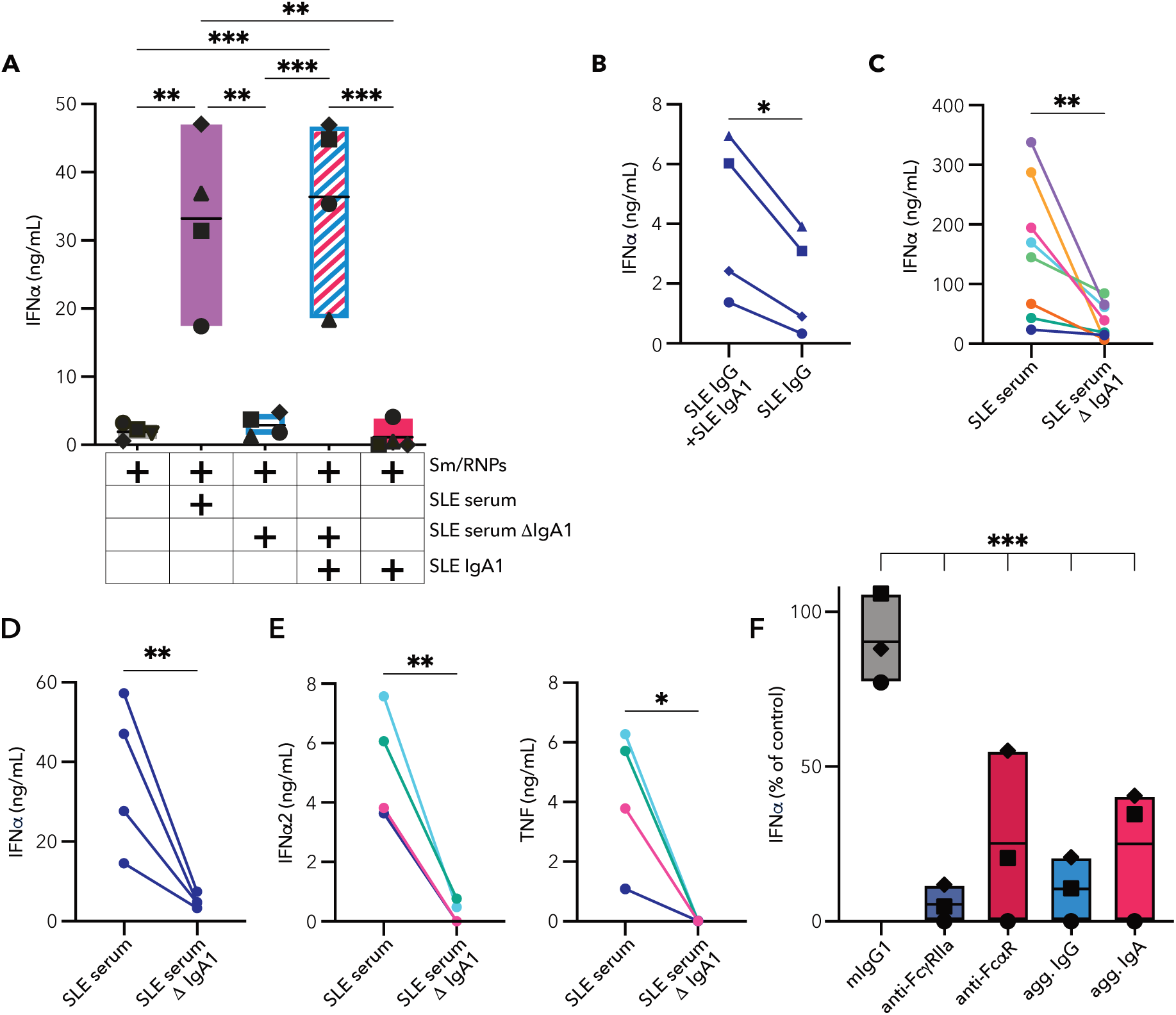
IgA1 in Sm/RNP immune complexes enhances IFNα production by pDCs. pDCs enriched from PBMCs freshly isolated from healthy control (HC) donor blood were incubated with immune complexes (ICs) generated with Sm/RNP RNA-containing nuclear antigens. **(A)** pDC IFNα secretion after incubation with Sm/RNPs alone or with Sm/RNP ICs generated with SLE serum, SLE serum ΔIgA1 or SLE IgA1. Each symbol shape represents a different pDC donor, mean and range shown. **(B)** Paired analysis of pDC IFNα secretion after incubation with Sm/RNP ICs generated with purified SLE IgG (135 μg) and purified SLE IgA1 (10 μg) or purified SLE IgG alone (135μg). **(C)** Paired analysis of pDC IFNα secretion after incubation with Sm/RNP ICs generated with either SLE serum or SLE serum ΔIgA1 from 8 SLE serum donors. **(D)** Paired analysis of pDC IFNα secretion after incubation with Sm/RNP ICs generated with either SLE serum or SLE serum ΔIgA1 from the same SLE serum donor and the same HC pDC donor assessed at 4 times over 14 months. **(E)** Paired analysis of pDC IFNα2 (*left*) and TNF (*right*) secretion after incubation with SLE serum or SLE serum ΔIgA1 ICs. **(F)** pDC IFNα secretion after pDCs were incubated with PBS, mIgG1 isotype control, anti-FcγRIIa, anti-FcαR, aggregated IgG or aggregated IgA for 2 hours prior to the addition ICs generated from SLE serum. Data are shown as a percent of PBS control with mean and range (raw data shown in Supplemental Figure 4E). (**A**) One way ANOVA, repeated measures, Bonferroni correction for multiple comparisons between all groups. Only significant comparisons are shown (*** p<0.001 and ** p<0.01). (**B-D**) Paired t-test (** p<0.01 and * p<0.05). **(F)** One way ANOVA, repeated measures, Bonferroni correction for comparing all blocking conditions to the isotype control (*** p<0.001). SLE serum donors; n=1 (A, B D and F), n=4 (E), n=8 (C). HC pDC donors; n=1 (C-E), n=3 (F), n=4 (A and B).

We corroborated the importance of IgA1 in pDC responses to Sm/RNP ICs with sera from an additional 7 SLE subjects, where we found a 38-99% reduction in IFNα secretion depending upon the serum donor (Fig. 4C, Supplementary Fig. 4B, Supplementary Table 3). We also validated the reproducibility and durability of these findings by repeating experiments using the same SLE serum donor with the same healthy control pDC donor different days months apart (Fig. 4D, Supplementary Fig. 4C). Consistent results were also found when using different healthy control pDC donors with the same SLE serum donor (Supplementary Fig. 4D). We detected low amounts of IFNβ secreted from 3 of 4 pDC donors when stimulated with SLE serum ICs containing both IgA1 and IgG. However, there was no detectable IFNβ in response to Sm/RNP ICs lacking IgA1 (Supplementary Fig. 4E). Measurement of IFNα2 specifically, as well as TNF, showed a similar strong dependence on IgA1 (Fig. 4E). Therefore, our data demonstrate a previously unreported role that IgA1 anti-nuclear antibodies have on IC activation of pDCs for cytokine secretion.

We hypothesized that Sm/RNP ICs containing both IgG and IgA1 would require co-engagement of Fc receptors for these isotypes, FcγRIIa and FcαR respectively, for potent IFNα production. To test whether FcγRIIa and FcαR are required for pDC recognition of Sm/RNP ICs, we blocked pDC FcγRIIa with aggregated purified IgG or anti-FcγRIIa monoclonal antibody (mAb), or blocked FcαR with aggregated purified IgA or anti-FcαR mAb before addition of ICs generated with SLE serum. IC-induced IFNα was completely dependent upon signaling through FcγRIIa, with >95% reduction in IFNα secretion with either aggregated IgG or 6C4 blocking. Consistent with the large effect we observed of IgA1 depletion on pDC responses to ICs, both anti-FcαR and aggregated IgA significantly reduced pDC IFN by ∼80% (Fig. 4F, Supplementary Fig. 4H). These data in combination with our IgA1-depletion results led us to conclude that responses to mixed ICs containing both IgA1 and IgG are mediated through co-engagement of the IgA-specific FcαR acting with the IgG-specific FcγRIIa.

### IgA1 in immune complexes promotes internalization by pDCs

We hypothesized that ICs that contain IgA1 might be able to better bind to and be internalized by pDCs, delivering the Sm/RNPs to endosomes where the RNA contained in the RNPs can activate TLR7. To visualize internalization of ICs, we labeled Sm/RNPs with AF647 and generated ICs that were detectable by flow cytometry and microscopy. Using flow cytometry, we found that IC binding and internalization peaked at ∼12 hours and by 20 hours detection of Sm/RNP-AF647 decreased presumably due to degradation inside the cells (Fig. 5A). When comparing the ICs generated with IgA1-sufficient SLE serum versus those generated with IgA1-depleted SLE serum, we found that at 6 and 12 hours there was less pDC binding and internalization of IgA1-depleted ICs (Fig. 5A). In subsequent experiments we assessed the 12-hour time point and found that binding and internalization was significantly decreased in IgA1-depleted Sm/RNP ICs compared to those generated with IgA1-sufficient SLE serum using pDCs from multiple HC donors (Supplementary Fig. 5A) and using ICs generated with serum from 4 SLE donors (Fig. 5B). Therefore, these data show IgA1 promotes binding and/or internalization of Sm/RNP ICs.

**Figure 5:**
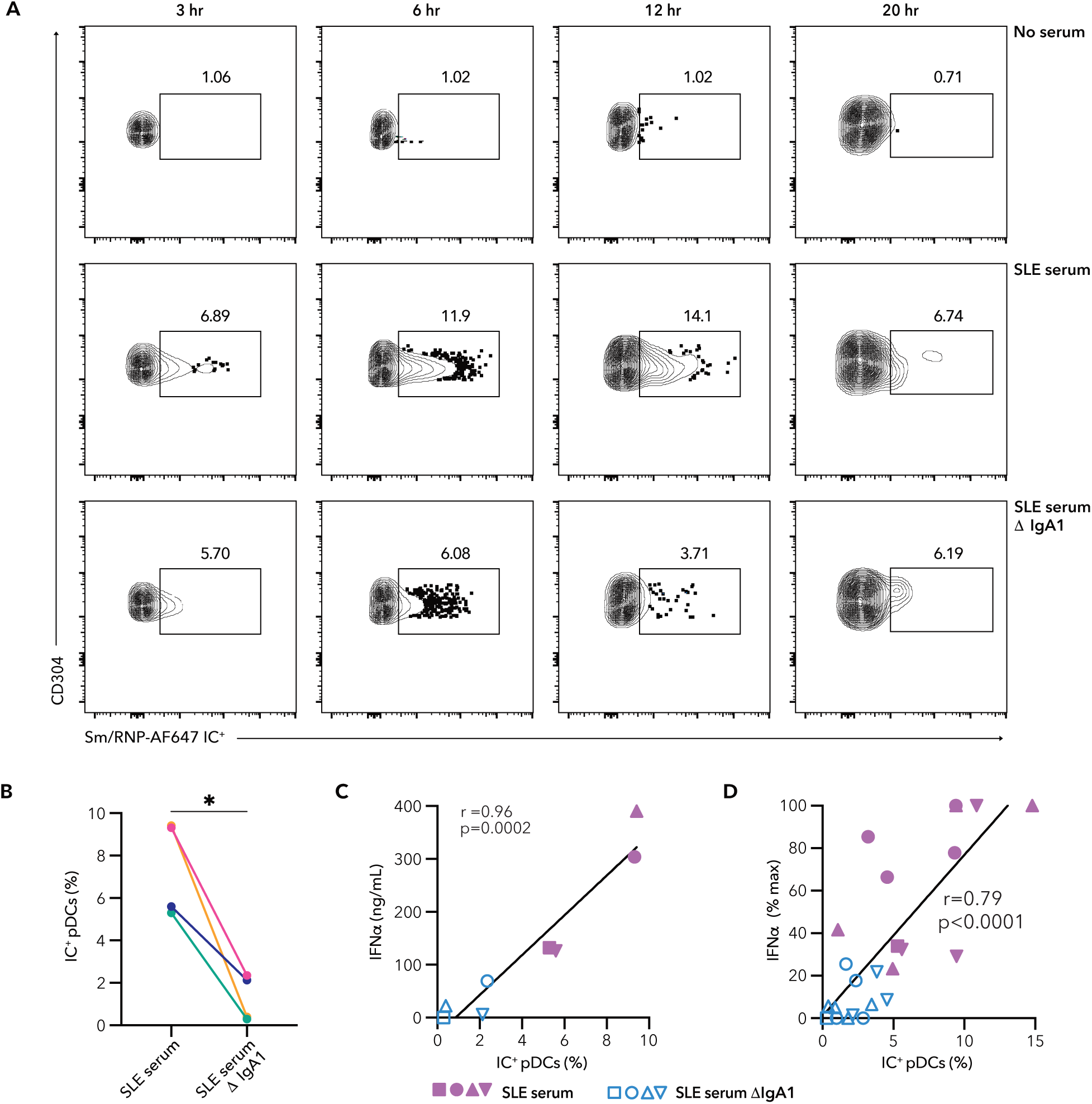
IgA1 autoantibodies contribute to IC association with pDCs. **(A)** pDCs were incubated with Sm/RNP-AF647 alone (*bottom row*) or Sm/RNP-AF647 ICs generated from SLE serum (*top row*) or SLE serum ΔIgA1 (*middle row*), for 3-20 hours. Representative flow cytometry plots show the percent of Sm/RNP-AF647^+^ pDCs. The Sm/RNP-AF647^+^ gate was determined by selecting where binding of Sm/RNP-AF647 control was approximately 1% or less (*bottom row*). **(B)** Percent IC^+^ pDCs at 12 hours after incubation with ICs generated with SLE serum or SLE serum ΔIgA1 from 4 SLE donors. **(C)** Correlation between IC^+^ pDCs (%) at 12 hrs and IFNα secretion at 20 hours (ng/mL) after incubation with Sm/RNP-AF647 ICs generated from SLE serum (*solid lavender symbols*) or SLE serum ΔIgA1 (*open blue symbols*). Plot shows the same IFNα secretion data from (B) with each symbol shape corresponding to the SLE serum donor. **(D)** Correlation of 12 hr IC^+^ pDCs (%) and normalized 20 hr IFNα production (% max) for combined experiments in which pDCs were incubated with Sm/RNP-AF647 ICs generated with SLE serum or SLE serum ΔIgA1. (**B**) Ratio paired t-test (* p<0.05) (**C, D**) r = Pearson’s correlation coefficient. SLE serum donors; n=1(A), n=4 (B, C, D). HC pDC donors; n=1 (A, B, C), n=4 (D).

When directly comparing the time course of IFNα production to corresponding Sm/RNP-AF647 internalization we found that there was detectable IFNα at 12 hours that was increased substantially at 20 hours (Supplementary Fig. 5B), while internalization peaked at 12 hours (Fig. 5A). We hypothesized that if increased internalization is the mechanism by which IgA1-IgG synergy occurs then Sm/RNP-AF647 internalization at 12 hours and IFNα production at 20 hours should correlate to one another. For this analysis we included both groups of ICs generated with either IgA1-sufficient or IgA1-depleted SLE serum. Consistent with our hypothesis, internalization of Sm/RNP ICs at 12 hours strongly correlated to IFNα secretion at 20 hours (r=0.96, p=0.0002) when assessed within a single experiment using one pDC donor and 4 SLE serum donors (Fig. 5C). When we combined data from multiple experiments significant correlation was maintained (r=0.66, p=0.0005) (Supplementary Fig. 5C) and this correlation was stronger (r=0.79, p<0.0001) when we normalized to account for the pDC donor differences in the magnitude of IFNα produced (Fig. 5D). Therefore, the degree of binding and internalization of Sm/RNP ICs correlated to the amount of IFNα produced by pDCs.

As we could not distinguish IC binding and IC internalization using the flow cytometry assay, we used confocal microscopy to assess IC internalization more directly. We added either IgA1-sufficient or IgA1-depleted AF647-labeled Sm/RNP ICs to freshly isolated HC pDCs and cultured for 12 hours, using Sm/RNP-AF647 alone as a control for non-specific uptake, and collected full z-stacks of imaged fields to determine whether the AF647 signal was fully internalized or bound to the pDC surface (Fig. 6A, Supplementary Fig. 6A, 6B). Using IgA1-sufficient SLE serum, we found that 29% of pDCs had internalized Sm/RNP-AF647 ICs and 9.6% had Sm/RNP-AF647 ICs that appeared bound to the surface of the cell (Fig. 6B). Sm/RNP-AF647 ICs made with SLE IgA1-depleted serum resulted in significantly reduced binding and internalization, with only 4.6% of pDCs showing internalization and 4.6% of pDCs showing binding of ICs. pDCs incubated with Sm/RNP-AF647 alone showed little binding or internalization (Fig. 6B). Taken together with our flow cytometry data, we conclude that IgA1 in ICs results in increased binding and internalization by pDCs that correlates with IFNα production.

**Figure 6:**
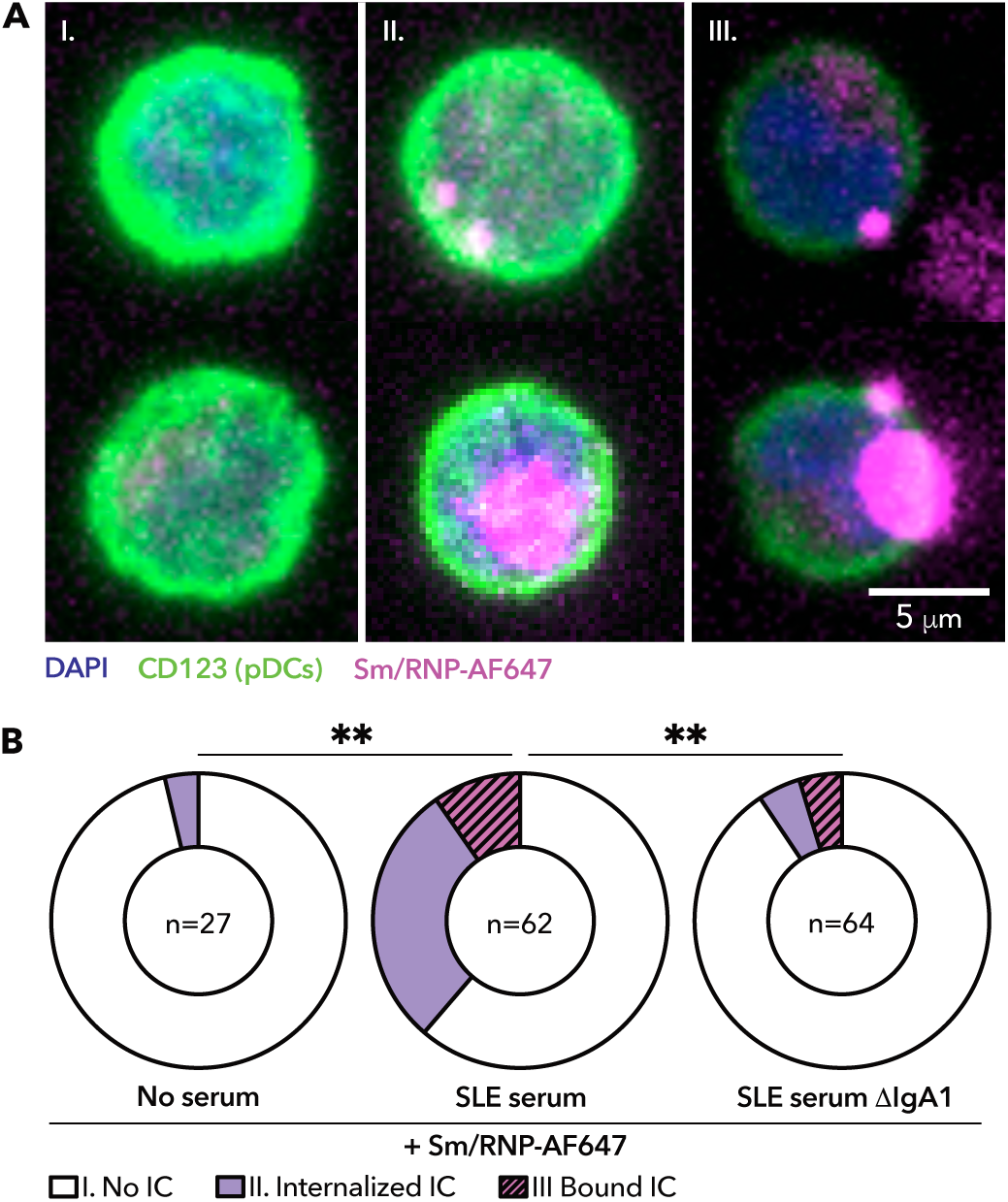
IgA1 autoantibodies enhance pDC internalization of immune complexes. Enriched pDCs from healthy individuals were incubated with Sm/RNP-AF647 alone or Sm/RNP-AF647 ICs generated with either SLE serum or SLE serum ΔIgA1 for 12 hrs (n=2 SLE serum donors). **(A)** Representative confocal images of the three categories of staining observed with DAPI (*blue*), CD123 (*green*) and Sm/RNP-AF647 ICs (*magenta*). Group I (*left*), pDCs that had no ICs bound or internalized or very faint IC signaling indicating low level or diffuse ICs bound. Group II (*middle*), pDCs with either small (*top cell*) or large (*bottom cell*) internalized ICs. Group III (*right*), pDCs with either small (*top cell*) or large (*bottom cell*) ICs that were bound but not internalized. Images shown are a composite of collapsed z-stacks, Supplemental Fig. 6 shows examples with full z-stacks to demonstrate the difference between internalized and bound ICs. **(B)** Donut plots show the proportion of pDCs in each group for each experimental condition: Sm/RNP-AF647 with no serum (*left*), Sm/RNP-AF647 ICs generated with SLE serum (*middle*), and Sm/RNP-AF647 ICs generated with SLE serum ΔIgA1 (*right*). The center of the donut plot shows the number of cells imaged and analyzed in each group. Chi-squared analysis performed between all groups and statistically significant comparisons shown (** p<0.01, ** p<0.01).

### SLE pDCs have increased capacity to bind ICs and increased surface FcαR expression compared to healthy control pDCs

Because IC internalization and pDC IFNα production correlated and both pDCs and IFNα are implicated in SLE pathology, we asked if pDCs from SLE donors had an increased capacity to bind to ICs compared to those from matched HC donors (Supplementary Table 3). We incubated PBMCs with Sm/RNP-AF647 ICs generated with SLE serum that contains both IgG and IgA for 3.5 hours at 37 degrees C and then measured binding and internalization by flow cytometry (Supplementary Fig. 7A). Importantly, the SLE serum used here showed strong IgA1 dependence for pDC IFNα secretion and IC binding/internalization in our assays. We found that SLE donor pDCs had significantly increased binding/internalization of Sm/RNP ICs compared to pDCs from HC donors (Fig. 7A). We also measured FcαR and FcγRIIa expression on duplicate samples from this cohort and found that pDC IC binding/internalization positively correlated to pDC FcαR surface expression, but not FcγRIIa surface expression (Fig. 7B). CD14^+^ monocytes in the same samples showed a trend for increased IC binding and internalization by those from SLE donors, but this did not reach statistical significance, nor did it correlate with monocyte FcαR or FcγRIIa expression (Supplementary Figs. 7B, 7C). Therefore, pDCs from donors with SLE had the capacity to bind and internalize IgA1-dependent ICs to a greater extent than those from HC donors.

**Figure 7.**
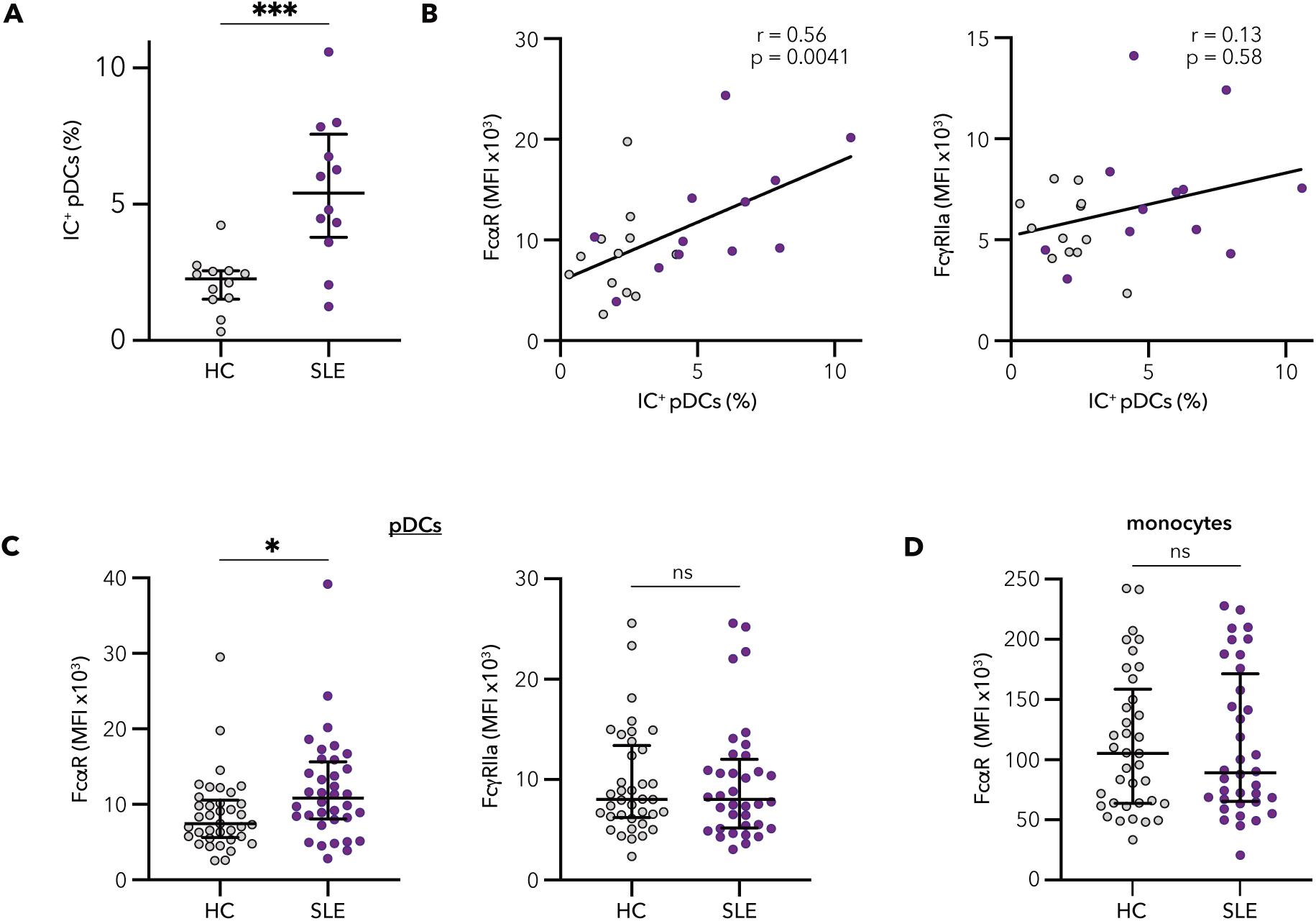
SLE pDCs have increased binding to immune complexes and increased expression of FcαR. **(A)** PBMCs from individuals with SLE (n=12) and matched healthy control subjects (n=12) were incubated for 3 hours with ICs generated from Sm/RNP-AF647 and SLE serum. Percent of gated pDCs with ICs is shown, determined as described in Figure 5A. **(B)** PBMCs from the same subjects as in **(A)** were also stained for surface FcαR and FcγRIIa and correlations between IC^+^ pDCs and FcαR (*left*) or FcγRIIa (*right*) are shown. Grey circles are pDCs from healthy control individuals and purple circles those from SLE individuals. (**C**) Cell surface expression of FcαR (*left*) and FcγRIIa (*right*) on gated pDCs from individuals with SLE (n=36) and matched HC subjects (n=37). **(D)** Cell surface expression of FcαR on gated monocytes from the same SLE and HC subjects in Fig. 7C. (**A**-**D**) Each circle represents an individual subject; (**A**, **C** and **D**) Student’s t-tests (* p<0.05 and ***p<0.001). (**B**) r=Pearson’s correlation coefficient.

Based on our findings that pDC IC binding/internalization and FcαR expression correlated, we hypothesized that the increased Sm/RNP IC internalization by SLE pDCs might be due to increased FcαR surface expression compared to HC pDCs. Therefore we used flow cytometry to measure expression of these receptors on pDCs from 36 matched SLE and HC donors (Supplementary Table 1, 3). FcαR expression was significantly higher on pDCs from SLE donors than from HC donors, while there was no difference between HC and SLE pDCs in FcγRIIa expression (Fig. 7C) or IgE specific FcγRIIa expression (Supplementary Fig. 7D). CD14^+^ monocytes from HC and SLE donors showed no difference in either FcαR or FcγRIIa expression (Fig. 7F, Supplementary Fig. 7E). These data suggest that the increased Sm/RNP IC binding/internalization by pDCs from SLE donors may be due to increased FcαR expression, which could lead to increased IFNα production from SLE pDCs in a feed forward loop.

To explore whether the expression of FcαR on SLE pDCs was related to IFNα production in vivo, we performed whole blood RNA-Seq on matched samples taken from the same blood draw that we had used to measure FcαR expression for 18 subjects with samples available. Upon analysis of the top 50 genes positively correlated with FcαR MFI, we noted many were known interferon-stimulated genes (ISGs) with 19 of these top 50 found in the Hallmark IFNα response gene set (Fig. 8A), though several others in this list are well known ISGs including *IFIT1, STAT1, OAS3* and *CCL2*. To explore this association between pDC FcαR expression and ISG signature more closely, we assessed if there was a correlation between the median expression of the entire Hallmark IFNα response gene set and pDC FcαR surface expression. There was a significant positive correlation between pDC FcαR expression and the Hallmark IFNα gene signature in SLE donors (Fig. 8B), while there was no such correlation with this gene signature and pDC FcγRIIa expression or monocyte FcαR or FcγRIIa expression (Fig. 8C, Supplemental Fig. 8A). Therefore, the IFNα rich environment in SLE specifically correlates with FcαR expression on pDCs.

**Figure 8:**
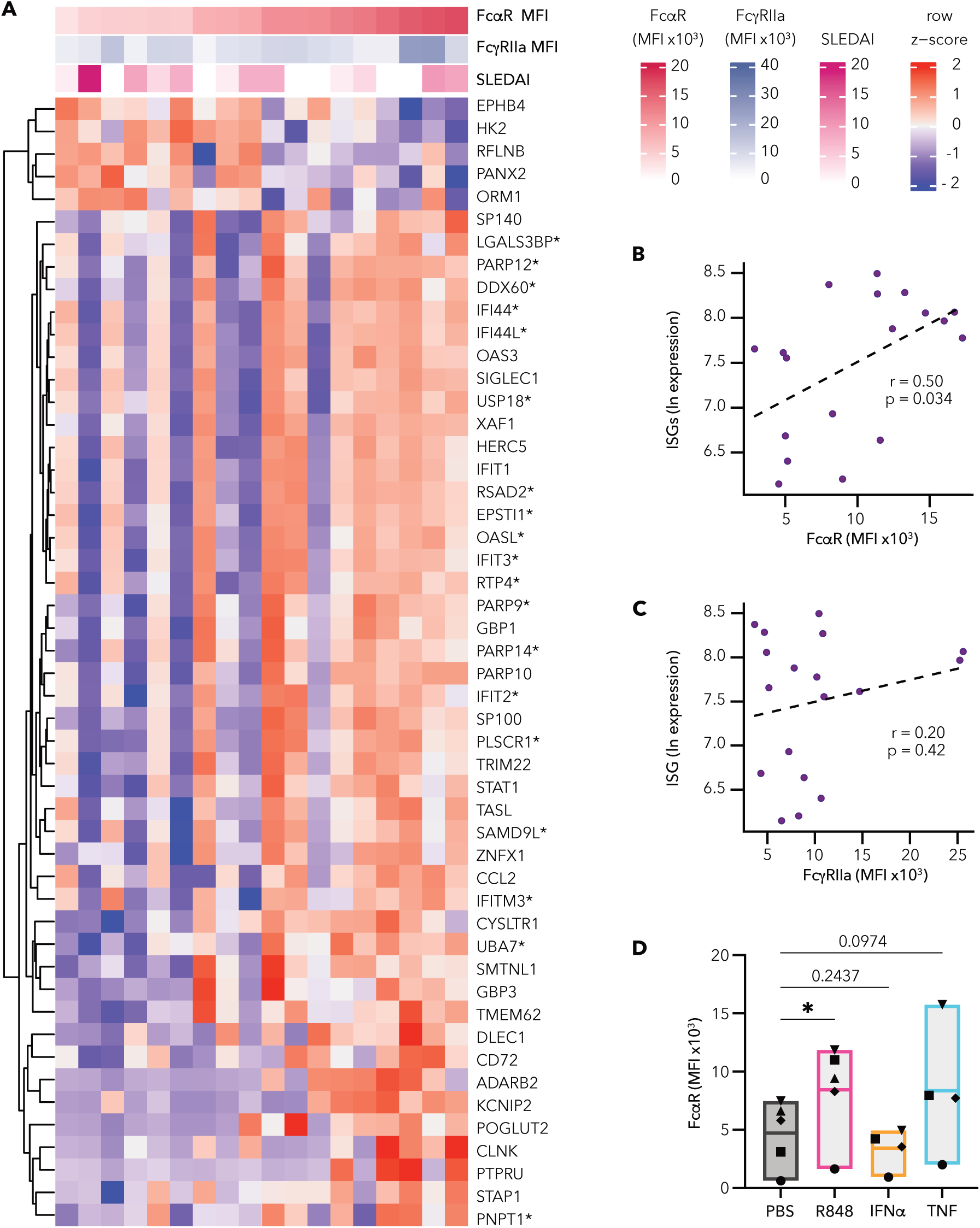
pDC FcαR expression correlates with IFN signature in SLE. **(A)** Whole blood RNA-Seq was performed for n=18 donors that were also assessed for FcαR and FcγRIIa expression (Fig. 7C). Heatmap shows the top 50 genes positively correlated to FcαR expression. Scales above the heatmap show FcαR MFI expression (red), FcγRIIa expression (blue) and SLEDAI score (pink). Genes that were found to be part of the Hallmark IFN-alpha response gene set from the Broad’s Molecular Signatures Database are indicated with an asterisk (*). **(B)** Correlation between cell surface expression of FcαR on gated pDCs and an interferon-stimulated gene (ISG) signature determined by RNA-seq performed on whole blood from 18 of the SLE subjects in (Fig. 7C). **(C)** Correlation between cell surface expression of FcγRIIa on gated pDCs and ISG signature determined by RNA-seq performed on whole blood from 18 of the SLE subjects in (Fig. 7C). **(D)** pDC FcαR expression (geometric MFI) after 36 hr incubation with PBS control, TLR7 agonist R848, IFNα or TNF. **(D)** Healthy control pDC donors n=4-5. (**B** and **C**) r = Pearson’s correlation coefficient. **(D)** One way ANOVA, repeated measures comparing all induction conditions to the PBS control (* p<0.05).

To examine if IFNα itself can upregulate pDC FcαR expression, we cultured pDCs for 36 hrs in the presence of IFNα and found there was no increase in pDC FcαR expression compared to untreated pDCs (Fig. 8D). We also investigated if treatment with the TLR7 agonist R848 or with TNF could upregulate surface FcαR on pDCs, and found that R848 treatment significantly upregulated pDC FcαR expression while there was a trend for increased pDC FcαR expression after TNF treatment (Fig. 8D). None of these treatments significantly changed FcγRIIa expression on pDCs (Supplementary Fig. 8B). Thus, while an ISG signature in SLE correlates with FcαR expression, this is likely due to pDC activation via TLR7, rather than a direct effect of type I IFNs on these cells.

## DISCUSSION

Our study aimed to better understand how IgA isotype autoantibodies contribute to SLE pathology. Our rationale for exploring IgA isotype autoantibodies in SLE was based upon recent insights including: the presence of IgA autoantibodies in SLE (*13–15, 17, 18, 47*), evidence that IgA can be pathogenic in rheumatoid arthritis (*48, 49*), and an increasing appreciation of the ability of IgA to facilitate proinflammatory immune responses when bound to viral and bacterial pathogens or in anti-tumor responses (*29, 30*). Glomerular IgA staining in addition to IgG, IgM, C1q and C3, so-called “full house” immunofluorescence, is an almost uniform finding in all classes of lupus nephritis (*16*). Additionally, IgA is the second most prevalent antibody in serum (*20*) and is produced at the highest rate of all isotype antibodies combined, with a rate of ∼66 mg/kg of body mass each day (*20*). Using autoantigen arrays, we found that most individuals with SLE have IgA autoantibodies against at least one nuclear antigen, and these individuals often have autoantibodies of both IgA and IgG isotypes against the same nuclear antigens. We also identified circulating immune complexes containing both IgA and IgG in some individuals with SLE, suggesting that increasing our knowledge of how these isotypes work together in immune complexes is essential for better understanding the mechanisms driving SLE pathology.

We found that IgA1 autoantibodies synergize with IgG to enhance pDC responses to immune complexes in SLE. When specifically investigating anti-Sm/RNP autoantibodies, we found that most SLE donors with anti-Sm/RNP IgG also had circulating IgA to this autoantigen. Using serum from these individuals, we identified a potent synergy between IgA1 and IgG anti-Sm/RNP autoantibodies for pDC cytokine production and pDC IC internalization. Because both FcαR and FcγRIIa signal through ITAMs, either in the associated FcRγ chain for FcαR or in the intracellular tail for FcγRIIa, it is not apparent why there is such strong synergy when co-signaling through these receptors when both IgG and IgA1 are present in ICs. A possible explanation for why IgA enhances the IC-mediated IFNα response is because the affinity for multimeric antigen-bound IgA for FcαR is greater than similarly complexed IgG for FcγRIIa (*50–52*). This could result in immune complexes containing both IgA1 and IgG having a higher overall avidity for pDC FcRs than those containing only IgG. Additionally, the FcRγ signaling chain associated with FcαR has two ITAMs, whereas FcγRIIa only has one ITAM (*28*), which may lead to stronger signaling for endocytosis of mixed IgA1-IgG ICs with twice the amount of ITAM signaling per IgA1 molecule bound than per IgG molecule bound. Perhaps unique, non-ITAM sequences in the FcαR cytoplasmic tail promote synergy. Furthermore, ICs that contain both IgA and IgG may increase internalization by pDCs because of the increased availability of Fc receptors for binding, whereas ICs that contain one isotype of autoantibody would only be able to bind FcαR or FcγRIIa, not both. The ability for an IC to bind to two different Fc receptors could be even more essential for a robust cytokine response when the surface expression of both receptors is relatively low, as is the case in pDCs, which we show express much less surface FcαR and FcγRIIa than monocytes from the same individual. Interestingly, platelets are also implicated in SLE pathogenesis with reported low-level FcαR and FcγRIIa expression (*53, 54*). Our data showing that IC binding and FcαR expression was greater on SLE pDCs compared to healthy controls also suggest that IgA1 enhancement of pDC IFNα production may be even more exaggerated in individuals with SLE than in healthy control individuals, exacerbating IFNα responses both to autoantigen-containing and viral-containing ICs in SLE.

IgA1 and IgG have different characteristics, which may result in qualitative changes in ICs containing both IgA1 and IgG autoantibodies compared to those without IgA1. IgA1 shares 90% sequence identity to IgA2 with the major difference being that IgA1 has a longer hinge region with no disulfide bonds allowing for increased flexibility compared to IgA2 and IgGs (*51, 55, 56*). While IgG3 also has longer hinge region, it retains the disulfide bonds (*57*). Interestingly one study found that longer hinge regions engineered into IgG1 and IgG3 antibodies resulted in increased phagocytosis in monocytes and neutrophils despite no change in affinity for the FcγRs (*58*). Even though this study focused on IgG hinge length and binding to FcγRs, their findings suggest that the increased hinge length of IgA1 could have a similar ability to increase internalization in pDCs via FcαR. Alternatively, the increased flexibility and length of the IgA1 hinge region may alter the size of ICs generated; larger ICs have been shown to have increased binding to FcRs (*59, 60*). In addition, large ICs that contain IgA and IgG could induce a higher degree of crosslinking of FcαR and FcγRIIa leading to stronger ITAM signaling and increased endocytosis. In our study, we did not determine the relative number or size of the ICs generated using IgA1-sufficient and IgA1-deficient SLE serum, and therefore we do not know if there were quantitative or qualitative differences in the ICs we generated. However, we found that Sm/RNP ICs generated with IgA1 alone were not sufficient to induce pDC IFNα secretion, though perhaps increasing the amounts of IgA1 used to generate ICs over the physiological amount found in SLE serum would result in pDC IFNα production with these IgA1-only ICs. The possibility remains that an excessive amount of IgA1 antibody could generate ICs capable of inducing pDC IFNα production, as IgA1 opsonized bacteria and viruses can induce phagocytosis in monocytes and phagocytosis and NETosis in neutrophils (*61–64*), though as discussed above the low FcαR and FcγRIIa expression on pDCs compared to these phagocytes may cause their responses to be regulated in a distinct manner. Despite these limitations, our data show a previously unrecognized connection between IgA1 and immune complex mediated pDC IFNα production and strongly suggests that IgA1 is a critical player in SLE.

As our immune system did not evolve to cause autoimmune diseases such as SLE, we speculate that this potent synergy between IgA and IgG in ICs evolved in the context of anti-viral immune responses, for which pDC IFNα production is protective (*65–67*). Perhaps this synergy evolved to allow for a strong, but transitory, pDC IFNα response to virus-containing ICs. While both IgA and IgG are produced during viral infection, IgA has a much shorter half-life (∼5 days) than IgG (∼21 days) (*68, 69*), which might allow for this synergy in pDC IFNα production to be important at the peak of the antibody response when viral antigens are present, but to resolve quickly as IgA antibodies with comparably short half-lives are removed from circulation. This may prevent maximal anti-viral IC IFNα production outside the peak of the response. In the case of SLE, self nuclear antigens are always present at low levels making IgA-IgG synergy maladaptive in the context of autoimmunity, and may promote pathogenic, chronic IFNα production. Increased nuclear antigens due to cell death during infection or injury or due to defective clearance of dying cells as is seen often in SLE may boost nuclear antigen-specific IgA production and subsequent pDC IFNα production, promoting disease flare in SLE. Thus, determining not only the presence of nuclear antigen-specific IgA, but also how it changes over time and may fluctuate with disease flare or severity is warranted. While we did not see a correlation between SLEDAI and IgA autoantibodies in our limited sample set, with larger cohorts there would be greater power to determine if the presence of IgA correlates to disease activity or any clinical manifestations of SLE.

The unique ability of FcαR to transduce either activating or inhibitory signals depending on whether the bound IgA is multimeric or monomeric, respectively (*25*), suggests that targeting FcαR therapeutically in SLE has two routes by which it could be beneficial. First, blocking FcαR could potentially prevent IgA-containing immune complexes from binding to and activating FcαR expressing cells, including pDCs, reducing IFNα production in response to ICs. If the blocking reagent is monomeric, it may also send inhibitory ITAM (ITAMi) signals through FcαR, which have only been shown in monocytes or macrophages to date (*26, 27*), but may also transduce similar signals in pDCs or neutrophils. In fact, one study has shown that early intervention targeting FcαR with a blocking mAb resulted in reduced disease outcomes in a pristane induced lupus nephritis model in transgenic mice expressing FcαR (*70*). While additional studies would be required to justify testing FcαR as a therapeutic strategy in SLE, they are hampered by the lack of an orthologue of FcαR in mice and therefore rely on expressing human FcαR as a transgene using promoters that faithfully mirror expression in human.

Our study has both strengths and limitations. Because we use ICs generated from SLE serum for all our studies, this has not allowed for us to define the anti-Sm/RNP IgA1 and IgG antibodies, their affinities and precise specificity, nor the optimal stoichiometry for generating pDC stimulatory ICs. We did not investigate how different IgG subtypes synergize with IgA1, nor did we isolate the effects of IgA2 subtype as most of serum IgA is IgA1 subtype (*44, 45*). We also did not examine the effects of other scarce isotype antibodies, such as IgD or IgE. It is important to note that Hennault et al. found that IgG and IgE anti-dsDNA antibodies could synergize to enhance pDC IFNα responses (*71*). For this reason, we measured total IgE and anti-SmRNP IgE in our reagents and found that the effects observed were not likely to be driven by IgE as there was no significant difference in total or anti-SmRNP IgE between serum and IgA-depleted serum (Supplementary Fig. 3A and 3C). Because we did not remove IgE from our reagents it is possible that all three isotypes synergize, and future studies will focus on better defining and isolating the contributions of each isotype antibody. However, the use of our less defined SLE serum instead of monoclonal antibodies ensures that we are observing biologically disease relevant phenomenon.

Additionally, isolating peripheral blood pDCs introduces donor variation and, due to the small amount of pDCs in blood, does not allow for high throughput testing of conditions optimal for inducing pDC IFNα responses to ICs. Using primary pDCs for our assays also did not allow for investigation of SLE pDC IFNα responses to ICs, as in SLE there is ∼50% reduction in number of pDCs and we obtain smaller amounts of blood from individuals with autoimmune disease. However, we were able to use SLE PBMCs to investigate pDC IC internalization and FcαR expression.

In conclusion, we show a remarkable ability of IgA1 isotype autoantibodies to synergize with IgG in RNA-containing ICs to generate a robust pDC IFNα response in both a FcαR and FcγRIIa dependent manner. We found that this pDC IFNα response correlated to pDC IC internalization and that SLE donor pDCs exhibited higher FcαR surface expression and more readily internalized IgA-containing ICs compared to HC pDCs. Taken together our data indicate that IgA1 ANAs are contributors in SLE pathology and warrant further investigation.

## MATERIALS AND METHODS

### Human samples

Frozen PBMCs, frozen serum, and fresh blood draws from HC and SLE subjects were obtained from the Benaroya Research Institute Immune-mediated Disease Registry and Repository. All pDC functional assays and microscopy imaging were performed with pDCs that were magnetically enriched from PBMCs isolated on the same day as the blood draw. Frozen SLE sera was used to generate ICs and for autoantigen arrays. See supplementary tables for specifics on donors used in each experiment. All experiments comparing cells from HC and SLE subjects were performed in a blinded manner. All experiments were approved by the Benaroya Research Institute Institutional Review Board.

### Kidney biopsy staining

Human kidney biopsy had standard pathologic workup, including light microscopic evaluation with Jones methenamine silver, periodic acid–Schiff, hematoxylin and eosin, and trichrome stains. For immunofluorescence (IF) microscopy, frozen tissue was stained with antibodies against IgG, IgA, IgM, C3, C1q, fibrin/fibrinogen, κ light chain, lambda light chain, and albumin.

### IgG and IgA Nuclear Antigen Microarray

Serum from 24 SLE subjects was assessed for IgG and IgA autoantigen binding at the University of Texas Southwestern (UTSW) Genomics and Microarray Core Facility. The autoantigen microarray super panel contained 128 autoantigens including most nuclear antigens relevant to SLE. For our analysis, we included 26 nuclear antigens against which SLE subjects commonly have antibodies. Normalized signal intensity (NSI) values were determined for each autoantigen as previously described by UTSW core researchers (*72, 73*). These NSI values were normalized to internal controls and corresponded to the amount of antibody binding to each autoantigen regardless of isotype, which allowed for direct comparisons between IgA and IgG. NSI values were then natural log (ln) transformed for correlation analysis and antibody heatmaps were created using the BioConductor package ComplexHeatmap. Because mean NSI values varied greatly between antigens, the heatmaps shown were also scaled for each antigen so that the maximum value was “1” and minimum value was “0”. Clustering of subjects and samples was conducted on the scaled IgA values, separately for RNA- and DNA-associated antigens, using Euclidean distances and complete linkage clustering as implemented in hclust; the clustering of IgA was then applied to the corresponding IgG heatmap. Supplementary Fig. 2B shows the raw mean NSI values.

### PBMC isolation and pDC enrichment from whole blood

240 mLs of freshly drawn whole blood was diluted 1:1 with PBS and underlaid with Ficoll-Paque plus (Cytiva). After centrifugation at 1000 x g for 25 minutes at 24 C without braking, PBMCs at the interface were collected, remaining RBCs were lysed with ACK Lysis buffer (Lonza), and subsequently washed 3 times with PBS 2% FBS (Sigma). pDCs were enriched from freshly isolated PBMCs using the human Plasmacytoid Dendritic Cell Isolation Kit II (Miltenyi Biotec) following manufacturer’s instructions with a 10% reduction in the recommended amounts of reagents. Enriched pDCs were counted and purity was measured by flow cytometry. pDC purity ranged from 60-99% (median 85%) with < 0.5% contamination from monocytes, B cells or T cells. For experiments using frozen PBMCs, sample vials containing 10-20 million PBMCs were thawed and washed in warmed complete RPMI (RPMI with 10% FBS, penicillin/streptomycin, L-glutamine). Complete RPMI for thawing cells also contained DNAse I (Sigma) at a concentration of 9 μg/mL.

### Staining for FcαR and FcγRIIa

Previously frozen PBMCs were resuspended in PBS and aliquoted into a 96 well plate for staining. All staining reagents used are listed in Supplementary Table 4. Cells were incubated with Zombie NIR live dead stain for 10 min at RT. After the live dead stain all subsequent washes and stains were performed in staining buffer (1% BSA, 2 mM EDTA in PBS) and plates were spun at 600 g for 2 min. After washing, cells were split into 3 replicate wells and incubated with purified anti-FcαR, anti-FcγRIIa or isotype control for 45 min at 4C. Samples were washed 3 times, then incubated with PE-labeled anti-mIgG1 secondary for 30 min at 4C. Samples were washed 3 times with staining buffer and once with 200 μg/mL mouse IgG1 to bind any residual anti-mIgG1 PE. Cells were then stained with directly conjugated antibodies to CD3, CD19, CD14, CD304, and CD123 for 25 minutes at 4C, washed and fixed in 3% paraformaldehyde. For experiments where samples were stained on multiple days, a replicate vial of control donor PBMCs was stained each day and MFI values of samples were normalized to the staining of control. All flow cytometry was performed on a Cytek Aurora spectral flow cytometer. Compensation/unmixing controls were done using single color stains with antibody at the same dilution used in the complete stain. Any sample in which <100 pDCs were collected were excluded from analysis. All flow cytometry analysis was done using FlowJo software.

### Staining for IgA and IgG

To generate aggregated IgA or IgG, biotinylated IgG (Novus NBP1-96855) or IgA (Novus NBP1-97181) was heated at a concentration of 2 mg/mL for 35-45 min at 65 C. After live dead staining, heat aggregated antibody was diluted in staining buffer to a final concentration of 0.5 mg/mL, incubated with PBMCs for 30 min at 37C. After washing cells were incubated with PE-labeled Streptavidin followed by washing, then staining for cell specific surface markers as above. To measure IgA and IgG bound to pDCs directly ex vivo, anti-IgA (AF647) and anti-IgG (FITC) were used.

### Anti-Sm/RNP and total antibody ELISAs

96 well flat bottom, high binding ELISA plates (Costar) were coated Sm/RNPs (Arotec Diagnostics ATR01) by adding 50 μl Sm/RNPs (0.5-1 μg/mL) diluted in PBS and incubated at 4 C overnight. Blocking was performed with 2X assay buffer (Invitrogen 88-50550-88) overnight at 4 C. Serum was serially diluted and incubated for 2 hrs at RT. After 7 washes with PBS/0.05% Tween-20, sample wells were incubated with biotin-labeled anti-IgA (Jackson Immunochemicals 309-065-008) or biotin-labeled anti-IgG (Jackson Immunochemicals 309-065-011) at 1:50,000 for 1 hr to detect IgA and IgG antibodies, respectively. Plates were washed as above and incubated with SA-HRP (eBiosciences 00-4100-94) for 30 mins followed by another wash cycle. TMB substrate was added, and reactions stopped with 0.16 M sulfuric acid once curves were detectable by eye (5-10 minutes). Then spectral reading was performed on VersaMax microplate reader (Molecular Devices). In addition to dilution curves, anti-Sm/RNP antigen binding was estimated in arbitrary units (AU) by using reagents from total IgG, IgA and IgE ELISA kits (ThermoFisher 88-50550-88, 88-50600-88, and 88-50610-88) to generate a standard curve for the appropriate isotype being measured using serial dilutions of IgA, IgG or IgE standards. anti-Sm/RNP isotype antibody signal was mapped on the corresponding anti-isotype antibody standard curve and scaled using arbitrary units. Total IgG, IgA and IgE were measured from serum using these same kits.

### IgA/IgG IC ELISA

To detect circulating ICs from donor serum, ELISA plates were coated with anti-human IgG (ThermoFisher 88-50550-88, 1X) and blocked overnight with 2X assay buffer (Invitrogen 88-50550-88) at 4C and for 2 hours at room temperature right before use. SLE donor serum was serially diluted in assay buffer, added to wells and then incubated for 2 hours at room temperature. After 7 washes with PBS/0.05% Tween-20, biotin anti-human IgA Fab (Invitrogen A24462) was added at a dilution of 1:100,000 for 2 hours at 4 C to detect human IgA. Plates were washed again, incubated with SA-HRP for 30 mins, washing and TMB substrate detection. Reactions were stopped with 0.16 M sulfuric acid. Spectral reading was performed on VersaMax microplate reader (Molecular Devices).

### IgA1 depletion from serum and IgA1 and IgG purification

IgA1 was depleted from SLE donor serum by affinity chromatography using Jacalin agarose (Pierce catalog #20395). SLE donor serum was first diluted 1:1 in PBS and an equal volume of Jacalin agarose resin was added to the column. Resin was added to 0.8 mL or 5 mL centrifuge columns (Pierce catalog #89868 or #89896) and washed 5 times with PBS. Diluted SLE serum was added to the columns and incubated for 1 hour at room temperature. Columns were spun at 2200 x g for 5 minutes to collect IgA-depleted serum, then were washed at least 8 times or until protein A280 measurement was below 0.05 mg/mL (Nanodrop spectrophotometer ND-1000). Two washes of 1M D-Galactose (Millipore Sigma PHR1206) were used to elute IgA1 from the column. Every 1 mL of collected IgA1 was concentrated by 10X using 10K molecular weight cut off protein concentrators (Pierce 88513). After initial concentration 10 additional washes with PBS at 10X remaining sample volume were performed in the concentrators to remove any remaining D-galactose. IgG was purified from IgA-depleted serum using Melon Gel IgG Spin Purification Kit (Thermofisher 45206) and following manufacturer’s instructions. Absorbance at 280 nm was measured to provide an estimated mg/mL amount of purified IgA1 or IgG in solution. Aliquots were frozen at -80 C for later use.

### Generation of Sm/RNP ICs

For each experimental well to be assayed, 12-16 μl of SLE donor serum or IgA1-depleted SLE serum was mixed with Sm/RNPs (Arotec ATR01) at 2-3 μg/mL in a total volume of 50 μl (diluted in PBS) and incubated for 1 hr at room temperature. Slightly different ratios of serum and Sm/RNPs were used depending on what was optimal for any given serum donor, however ratios were kept identical for all SLE serum and ΔIgA1 serum comparisons within a given donor. In some cases, ICs were made with Alexa Fluor 647 (AF647)-labeled Sm/RNPs. 0.5 mls of Sm/RNPs (0.75 mg/mL) were labeled using the Alexa Fluor 647 Protein Labeling Kit (Invitrogen A20173) following the manufacturer’s instructions without removal of unbound AF647. SmRNP-ICs were also generated with IgG purified from IgA1-depleted serum with or without the addition of purified SLE IgA1. 135 μg of SLE IgG either alone or mixed with 10 μg of purified IgA1 from the same SLE serum donor was incubated with Sm/RNPs in a total volume of 50 μl.

### IC stimulation of pDCs

25,000 – 45,000 pDCs were added to 96 well U bottom plate wells in 150 μl of culture medium (RPMI with 2% heat inactivated human AB serum (Omega Scientific HS-20), 20 ng/mL IL-3 (Peprotech 200-03), 200 U/L penicillin and 200 μg/L streptomycin). Cells were rested at 37°C for 1-3 hours before adding 50 μl of Sm/RNP ICs. After the addition of the ICs, plates were incubated for 20-24 hrs then 100 μl of supernatants were removed and stored at 4C. For blocking experiments, 25 μl of blocking reagents, isotype control antibody, or PBS were added to wells 2 hours before the addition of Sm/RNPs ICs. Blocking reagents were left in wells throughout the assay. Heat aggregated IgA (Invitrogen 31148) and IgG (Invitrogen UJ2861463) stock concentrations were 2 mg/mL; anti-FcαR MIP8a (Invitrogen MA5-28106), anti-FcγRIIa 6C4 (Invitrogen 16-0329-85) monoclonal and isotype control (Invitrogen 16-4714-85) stock concentrations were 1 mg/mL.

### Cytokine measurements from supernatants

HeLa cells stably expressing an ISRE (interferon stimulated response element) luciferase reporter construct were a generous gift from Dr. Dan Stetson (University of Washington) (*74*) and were used to measure type I IFNs in supernatants. 60,000 HeLa-ISRE cells were added to each well of 96 well white opaque tissue culture plates in a volume of 100 μl and cultured overnight. IFN Supernatants were serially diluted in the culture medium and 50 μl added per well. *E. coli* expressed human IFNα 2a (PBL Assay Science 11101) was serially diluted 1:2 to generate a standard curve from 1 mg/mL to 1.9 μg/mL. Plates were incubated at 37 C for 5-6 hours. After incubation, 50 μl of Bright-Glo Luciferase reagent (Promega E2620) was added to each well. Luminescence was measured using a SpectraMax iD3 (Molecular Devices). In some cases, cytokines in supernatants were measured using Bioplex assays (BioLegend) measuring IFNα-2 (catalog #740351), IFNβ (catalog #740353), and TNF (catalog#740359) according to manufacturer’s protocols. HeLa ISRE cells detect both IFNα and IFNβ. However, because the amount of IFNβ measured was over 1000 fold less than the total type 1 IFNs measured by our ISRE reporter assay used throughout the manuscript, we labeled the y-axis in graphs using this assay as IFNα.

### Immune complex internalization microscopy

Freshly enriched pDCs from healthy individuals were incubated Sm/RNP-AF647 alone or with ICs generated with Sm/RNP-AF647 and SLE serum or SLE serum ΔIgA1 for 12 hrs, and then stained with anti-CD123 to identify pDCs. pDCs were mounted to glass slides using ProLong™ Diamond Antifade Mountant with DAPI (Thermofisher P36962) and imaged on Leica SP5 confocal microscope. Multiple Z-stacked images were taken per cell and images processed using Image J. pDCs were then blindly categorized into three groups (no Sm/RNP-AF647 ICs, internalized Sm/RNP-AF647 ICs or surface bound Sm/RNP-AF647 ICs).

### Interferon Stimulated Genes (ISGs) measurement from whole blood

Whole blood was collected in Tempus Blood RNA Tubes (ThermoFisher), and RNA was extracted using MagMax for Stabilized Blood Tubes RNA Isolation Kit, followed by globin reduction using GlobinClear Human (ThermoFisher). To generate sequencing libraries, total RNA (0.5 ng) was added to reaction buffer from the SMART-Seq v4 Ultra Low Input RNA Kit for Sequencing (Takara), and reverse transcription was performed followed by PCR amplification to generate full length amplified cDNA. Sequencing libraries were constructed using the NexteraXT DNA sample preparation kit (Illumina) to generate Illumina-compatible barcoded libraries. Libraries were pooled and quantified using a Qubit® Fluorometer (Life Technologies). Sequencing of pooled libraries was carried out on a NextSeq 2000 sequencer (Illumina) with paired-end 59-base reads, using a NextSeq P2 sequencing kit (Illumina) with a target depth of 5 million reads per sample. Base calls were processed to FASTQs on BaseSpace (Illumina), and a base call quality-trimming step was applied to remove low-confidence base calls from the ends of reads. The FASTQs were aligned to the GRCh38 human reference genome, using STAR v.2.4.2a, and gene counts were generated using htseq-count (*75*). Quality metrics were calculated using the Picard family of tools (v1.134; http://broadinstitute.github.io/picard). We used quality thresholds of at least 2.5M total reads, at least 70% of reads aligned to human genome, and median CV of coverage at most 0.7; 18 out of 19 samples passed these filters and were included in subsequent analyses. Gene counts were filtered to only protein-coding genes with at least one count per million in 10% of libraries and normalized using the trimmed mean of m-values (*76*). Differential gene expression was determined using limma-voom, with FcαR MFI on pDCs as a continuous variable. We quantified ISG expression using the median log2 normalized counts of gene in the MSigDB Hallmark IFNα response gene set. The gene expression heatmap was created using ComplexHeatmap.

### pDC induction and measurement of FcαR

pDCs were enriched and cultured as previously described. PBS, R848 (2.5 μg/mL), IFNα (20 ng/mL) or TNF (20 ng/mL) were added to wells. After 36 hrs of culture, FcαR and FcγRIIa staining were performed as previously described.

### Statistics

Statistics and tests used are described in the figure legends and methods or results for each section. Calculations were performed using GraphPad Prism and R. Additionally, because Pearson correlations can be driven by outliers, we also calculated Spearman rank correlations and found qualitatively similar results.

## General

We thank Drs. Susan Canny, Christian Lood, Adam Lacy-Hulbert, Keith Elkon, David Rawlings, Ram Savan and Jim Zimring for input on this work, Dr. Anne Hocking and Taylor Lawson for input and editing the manuscript, and members of the Hamerman laboratory for input throughout this project. We thank Griff Gessay, Fanny Vaca Flores and Noah Biru for assistance with blood processing, Drs. Hannah Volkman and Dan Stetson for ISRE-luciferase reporter cells and Dr. Caroline Stefani for assistance with confocal microscopy. We also thank members of the Clinical, Cell and Tissue Analysis, and Genomics Cores at the Benaroya Research Institute and all blood donors.

## Author Contributions

Conceptualization: HRW, JAH, JHB Data Curation: MJD, SP Formal Analysis: MJD, HRW Methodology: HRW, MJD Investigation: HRW, CZ, SP, MN, LZL Visualization: HRW, MJD, SP, KDS Funding acquisition: JAH, HRW Supervision: JAH Writing – original draft: HRW, JAH, MJD Writing – review & editing: all authors

## Data, code and materials availability

All data are available in the main text or the supplementary materials, except for whole blood RNA-Seq data which is deposited NCBI GEO repository, accession number GSE242721. Custom code for analysis and visualization of the autoantigen microarray and RNA-seq data is available at https://github.com/BenaroyaResearch/Waterman_Hamerman_SLE_autoantibodies_pDCs_2024.

## SUPPLEMENTARY MATERIALS

**Supplementary Figure 1:**
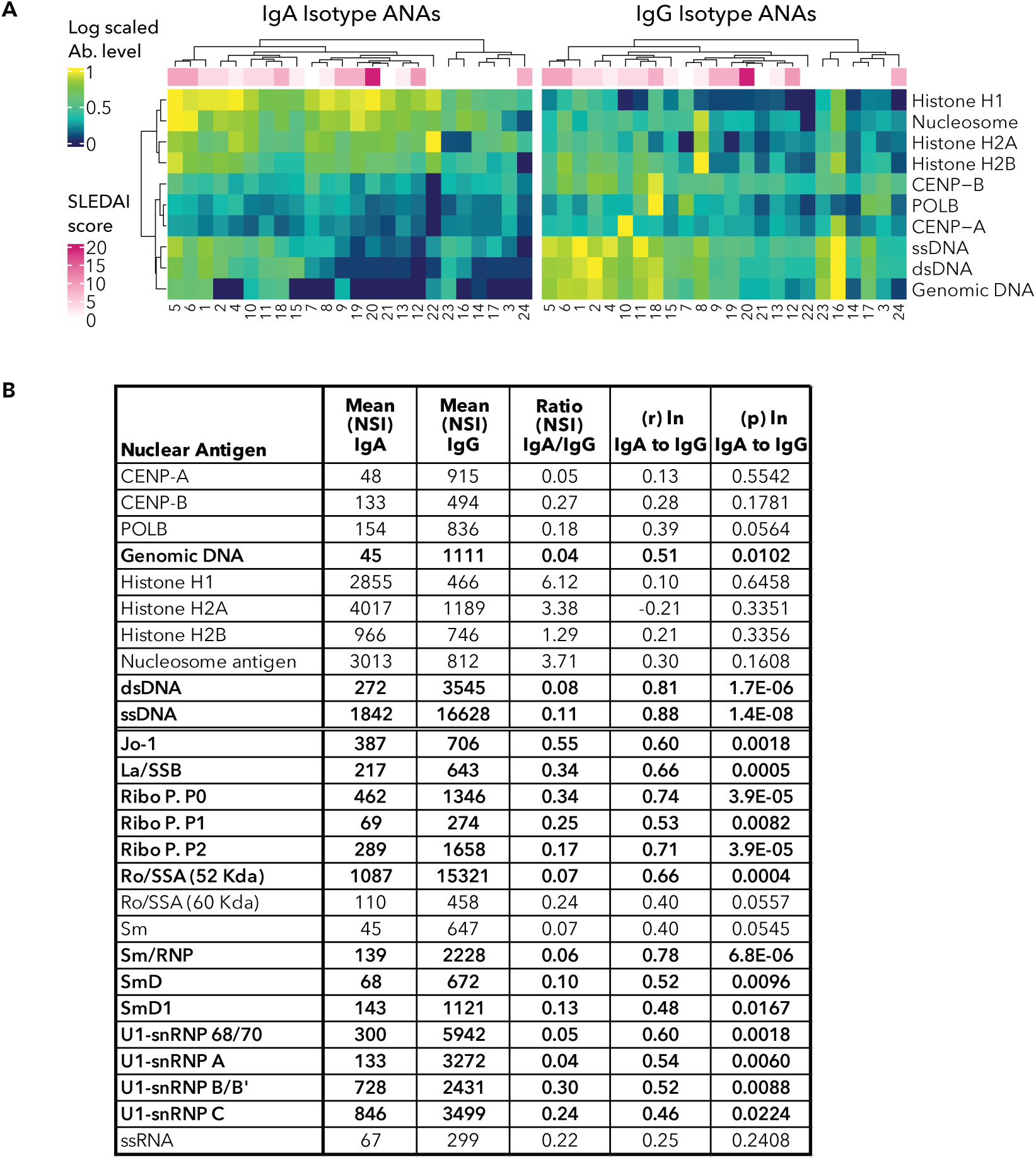
Supplementary figures relating to main Figure 1. **(A)** Autoantibody profiling of serum from 24 SLE subjects. Heatmaps of log-scaled antibody binding for IgA (*left*) and IgG (*right*) isotype antibodies to 10 DNA-associated nuclear antigens. Each row corresponds to an antigen and each column corresponds to an individual subject. The numbering of subjects below heatmap correspond to the number in Fig. 1A. The color bar across the top indicates SLEDAI score for each donor at time of blood draw. Clustering of subjects and samples is based on similarity in scaled IgA levels, with the same clustering applied to the IgG heatmap. **(B)** Table shows mean normalized signal intensity (NSI) values for IgA and IgG isotype antibody, the ratio of IgA/IgG NSI values, as well the Pearson’s correlation and p values for natural log transformed IgA and IgG NSI values.

**Supplementary Figure 2:**
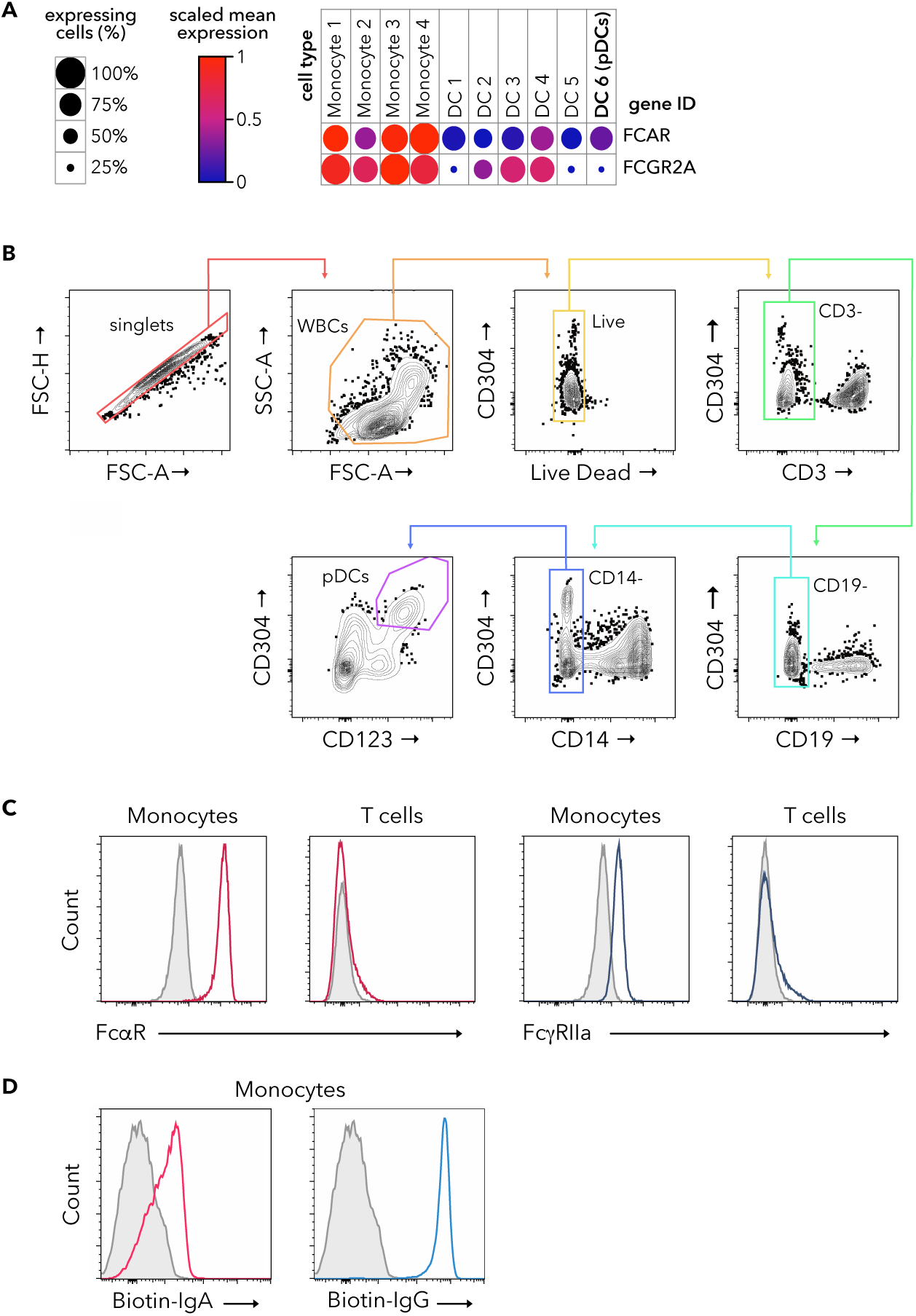
Supplementary figures relating to main Figure 2. **(A)** *FCAR* and *FCGRIIA* mRNA expression in monocytes (1–4) and DC (1–6) subsets analyzed from publicly available single cell RNA sequencing data (*43*). Color of the circles represent scaled mean expression levels and size of the circle corresponds to the percent of expressing cells for each group. **(B)** For all flow cytometry experiments gating for pDCs was done as shown. pDCs were identified as CD3^-^ CD19^-^CD14^-^CD304^+^CD123^+^ live single cells (*purple gate*). Monocyte and T cells were identified as CD3^-^ CD19^-^CD14^+^ and CD19^-^CD14^-^CD3^+^ live single cells respectively. **(C)** PBMCs were analyzed by flow cytometry for surface FcαR (*red*) and FcγRIIa (*blue)* staining with isotype control staining in grey. The first two histograms show FcαR staining on gated monocytes (*first panel*) and T cells (*second panel*). The last two histograms show FcγRIIa on gated monocytes (*third panel*) and T cells (*fourth panel*) **(D)** Flow cytometry analysis showing aggregated biotinylated IgA (*red, left*) and IgG (*blue, right*) staining on monocytes as detected by labeled streptavidin. Gray histogram shows control streptavidin staining alone.

**Supplementary Figure 3:**
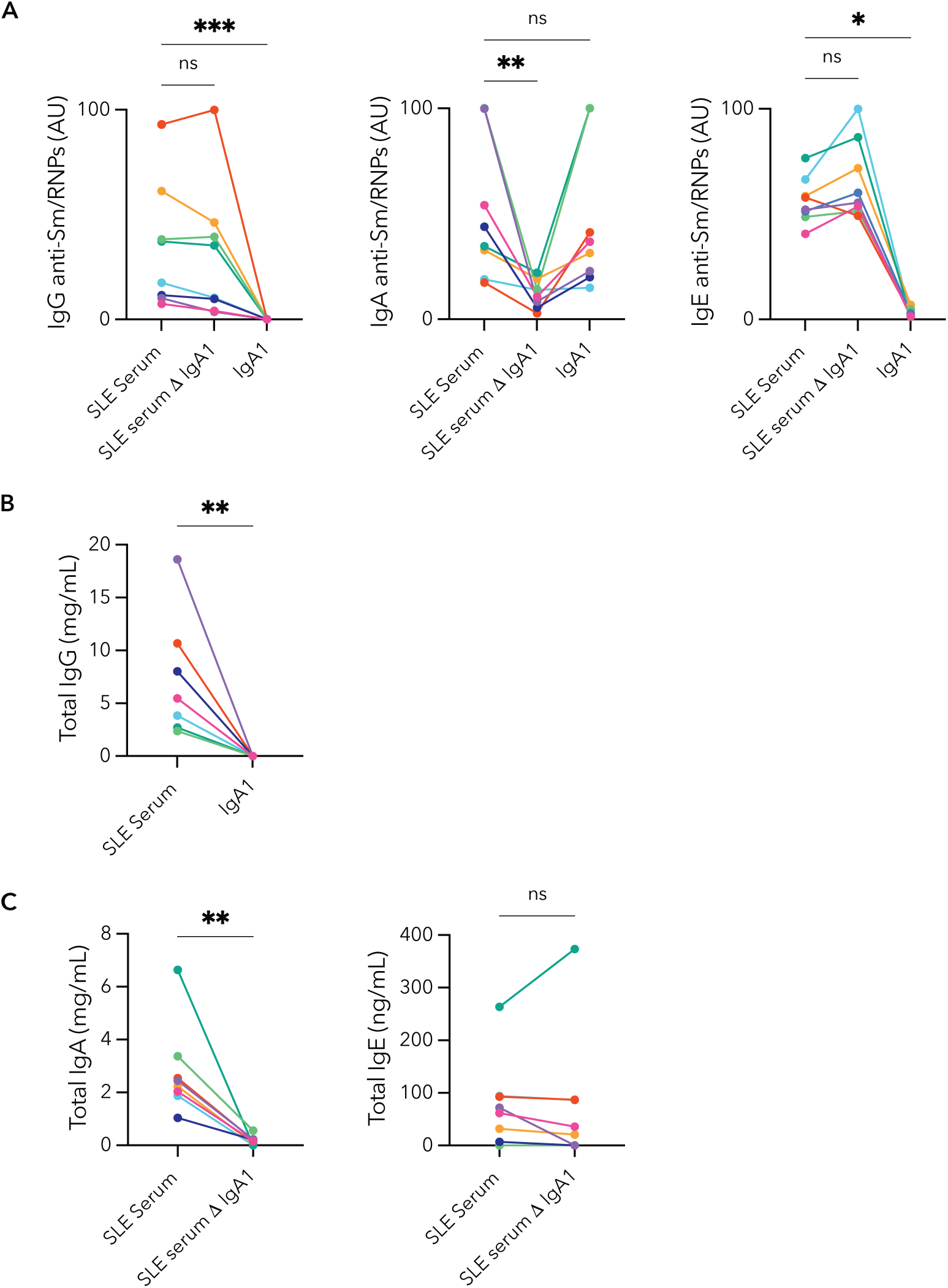
Supplementary figures relating to main Figure 3. **(A)** IgG (*left*), IgA (*middle*) and IgE isotype (*right*) anti-Sm/RNP was measured in arbitrary units (AU) as described in methods. SLE serum, SLE serum ΔIgA1 and purified IgA1 were assayed for the 8 SLE serum donors used for all pDC IFNα production experiments. Each donor is color coded the same throughout this figure and in all other figures. **(B)** Total IgG in SLE serum and SLE IgA1. **(C)** Total IgA (*left*) in SLE serum and SLE serum ΔIgA1. Total IgE (*right*) in SLE serum and SLE serum ΔIgA1.

**Supplementary Figure 4:**
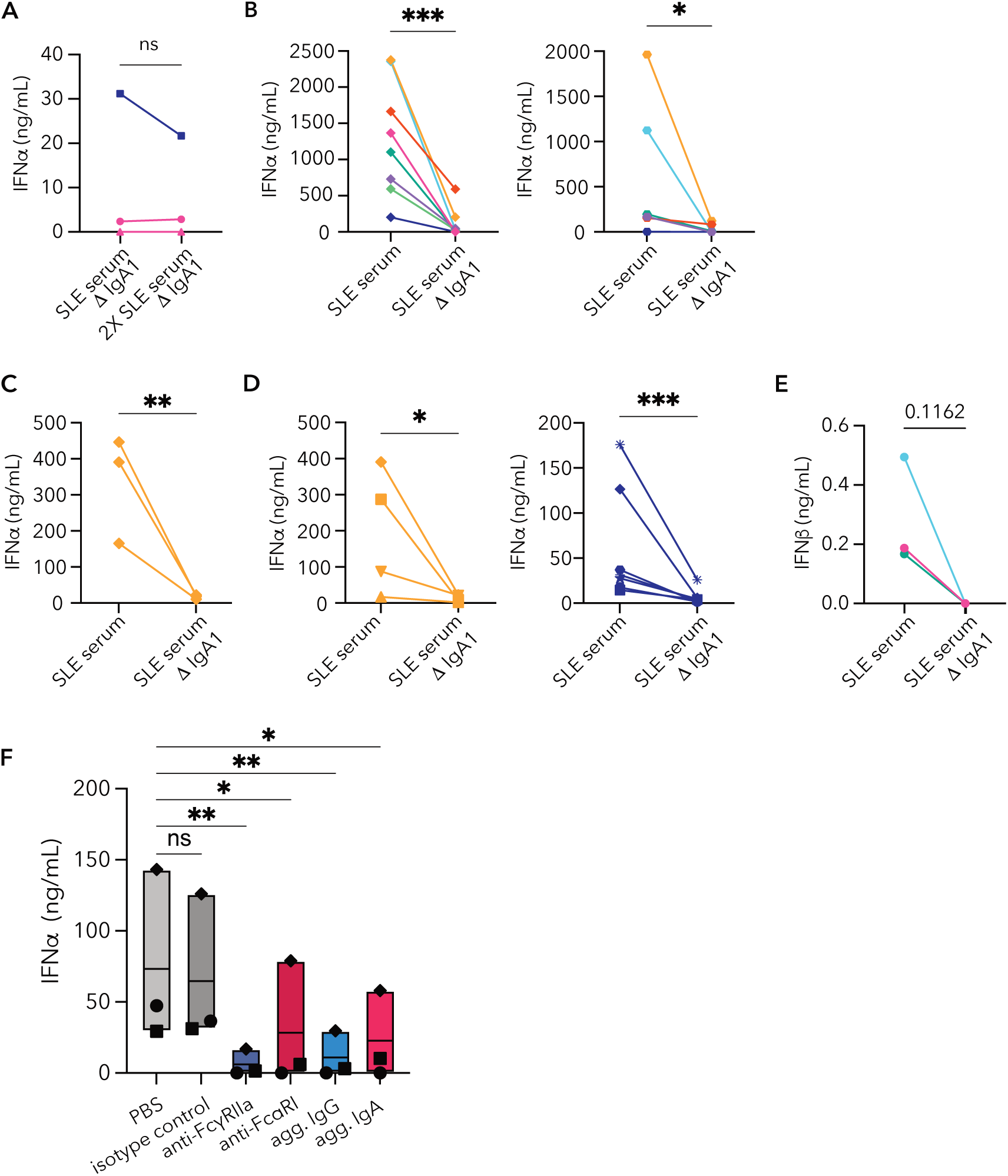
Supplementary figures relating to main Figure 4. pDCs enriched from PBMCs freshly isolated from healthy control (HC) donor blood were incubated with immune complexes (ICs) generated with Sm/RNP RNA-containing nuclear antigens. **(A)** IFNα produced by pDCs after 24 hr incubation with Sm/RNP ICs generated with either SLE serum ΔIgA1 or twice the amount of SLE ΔIgA1 serum (2X SLE ΔIgA1 serum). **(B)** pDC IFNα production after 30-hour incubation with Sm/RNP ICs made with SLE serum or SLE serum ΔIgA1 from multiple SLE donors. Each line represents a distinct SLE serum donor, and each panel is from a different HC pDC donor distinct from the donor in Fig. 4C. **(C)** Paired analysis of pDC IFNα secretion with the same SLE serum donor and HC pDC donor assessed at 3 times over 7 months. **(D)** HC pDCs from multiple donors were stimulated with Sm/RNP ICs made with SLE serum or SLE serum ΔIgA1 from one SLE serum donor (*left* same SLE serum donor as in Supplementary Fig. 4C, *right* same SLE donor from Fig. 4C). **(E)** Paired analysis of pDC IFNβ secretion after incubation with SLE serum or SLE serum ΔIgA1 ICs **(F)** Raw non-normalized data for experiment shown in Fig. 4F. HC pDCs were blocked with either PBS, isotype control antibody, monoclonal anti-FcγRIIa (6C4), monoclonal anti-FcαR (MIP8a), aggregated IgG or aggregated IgA for two hours before given Sm/RNP ICs generated with SLE serum. One way ANOVA with repeated measures (p = 0.018) with Bonferroni correction for multiple comparisons. Statistics for A, B and D-G are ratio paired t tests (* p<0.05, ** p<0.01, *** p<0.001.) SLE serum donors n=1 (C, D, F), n=2 (A), n=3 (E) n=6 (B *left*), n=8 (B *right*). HC pDC donors n=1 (B, C), n=3 (A, F), n=4 (D *left*), n=7 (D *right*).

**Supplementary Figure 5:**
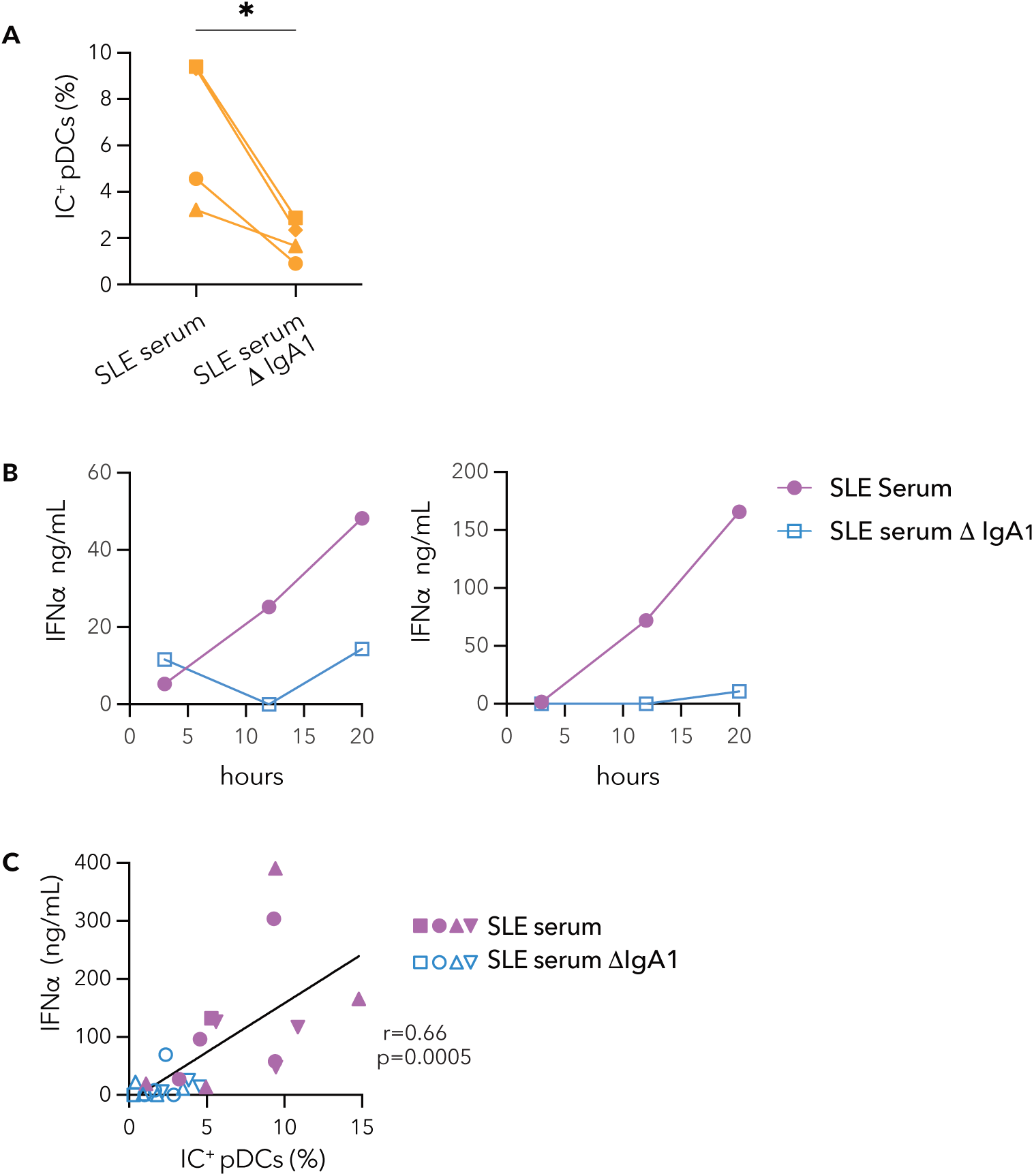
Supplementary figures relating to main Figure 5. ICs were generated AlexaFluor647-labeled Sm/RNPs (Sm/RNP-AF647). **(A)** Paired analysis of percent IC^+^ pDCs (n=4) after incubation with Sm/RNP-AF647 ICs generated with SLE serum or SLE serum ΔIgA1 from 1 SLE serum donor. **(B)** pDCs (n=1) were incubated with Sm/RNP-AF647 ICs generated with (n=2) SLE serum or SLE serum ΔIgA1 and IFNα secretion from replicate wells was measured at 3, 12 and 20 hours. Each graph shows data from a different SLE serum donor. **(C)** Correlation of 12 hour IC^+^ pDCs (%) and 20 hour IFNα production for combined experiments in which pDCs were incubated with ICs generated with Sm/RNP-AF647 mixed with either SLE serum or SLE serum ΔIgA1. The combined data were generated with pDCs from n=4 HC pDC donors and n=4 SLE serum donors. r = Pearson’s correlation coefficient.

**Supplementary Figure 6:**
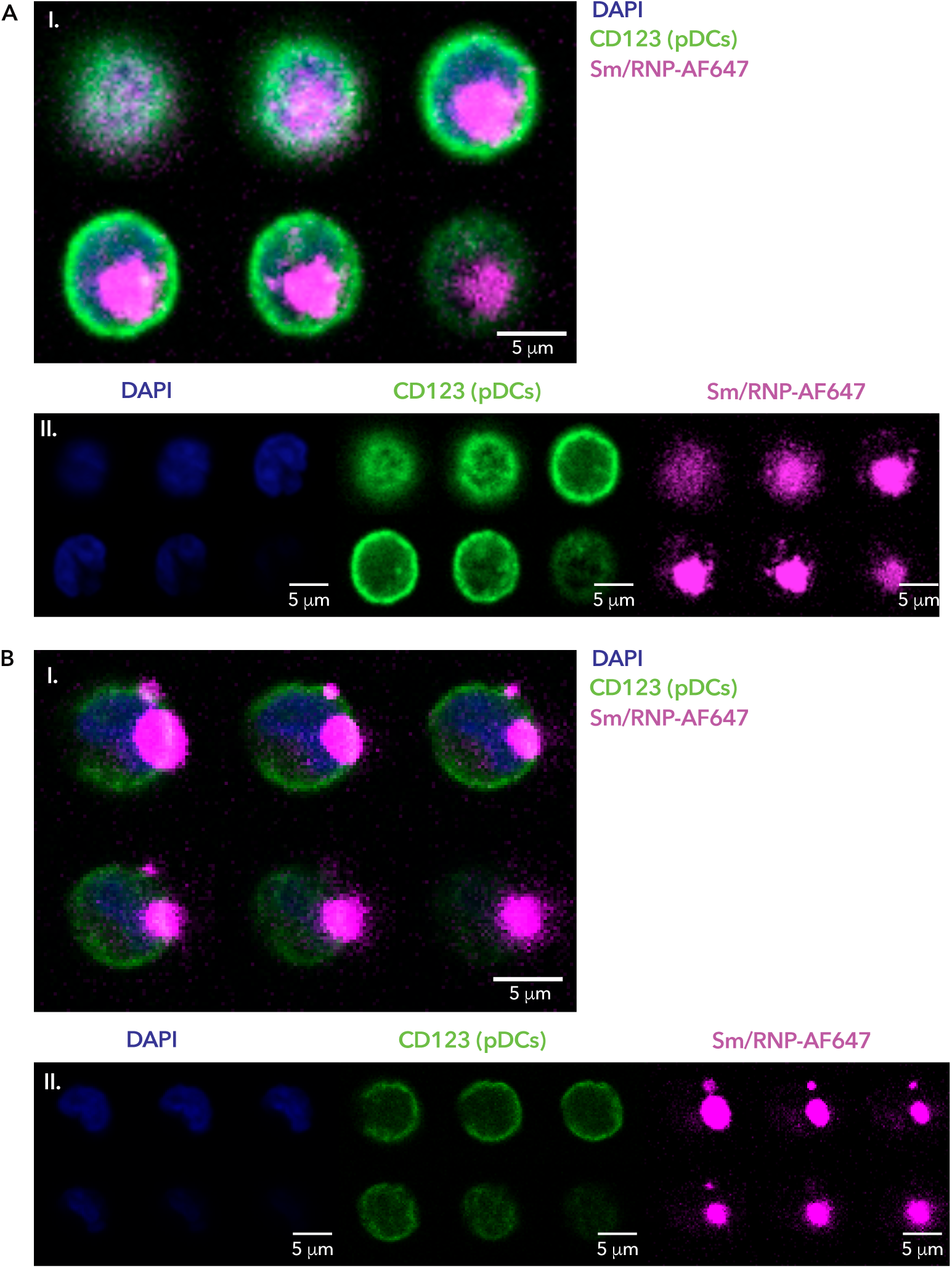
Representative images used to categorize Sm/RNP-AF647 SLE serum IC internalization and binding to pDCs. Healthy control (HC) pDCs were incubated with ICs generated with Sm/RNP-AF647 alone or Sm/RNP-AF647 with SLE serum or SLE serum ΔIgA1 for 12 hours as in described in Fig. 6. **(A)** Representative example of a cell with large, internalized ICs. **(B)** Representative example of a cell with large ICs bound to the outside of the cell. For both **(A)** and **(B)** the first series of images (I.) shows the z-stack with all channels and second series of images (II.) show the z-stack for each individual channel; DAPI (*blue*), CD123 (*green*) and Sm/RNP-AF647 ICs (*magenta*).

**Supplementary Figure 7:**
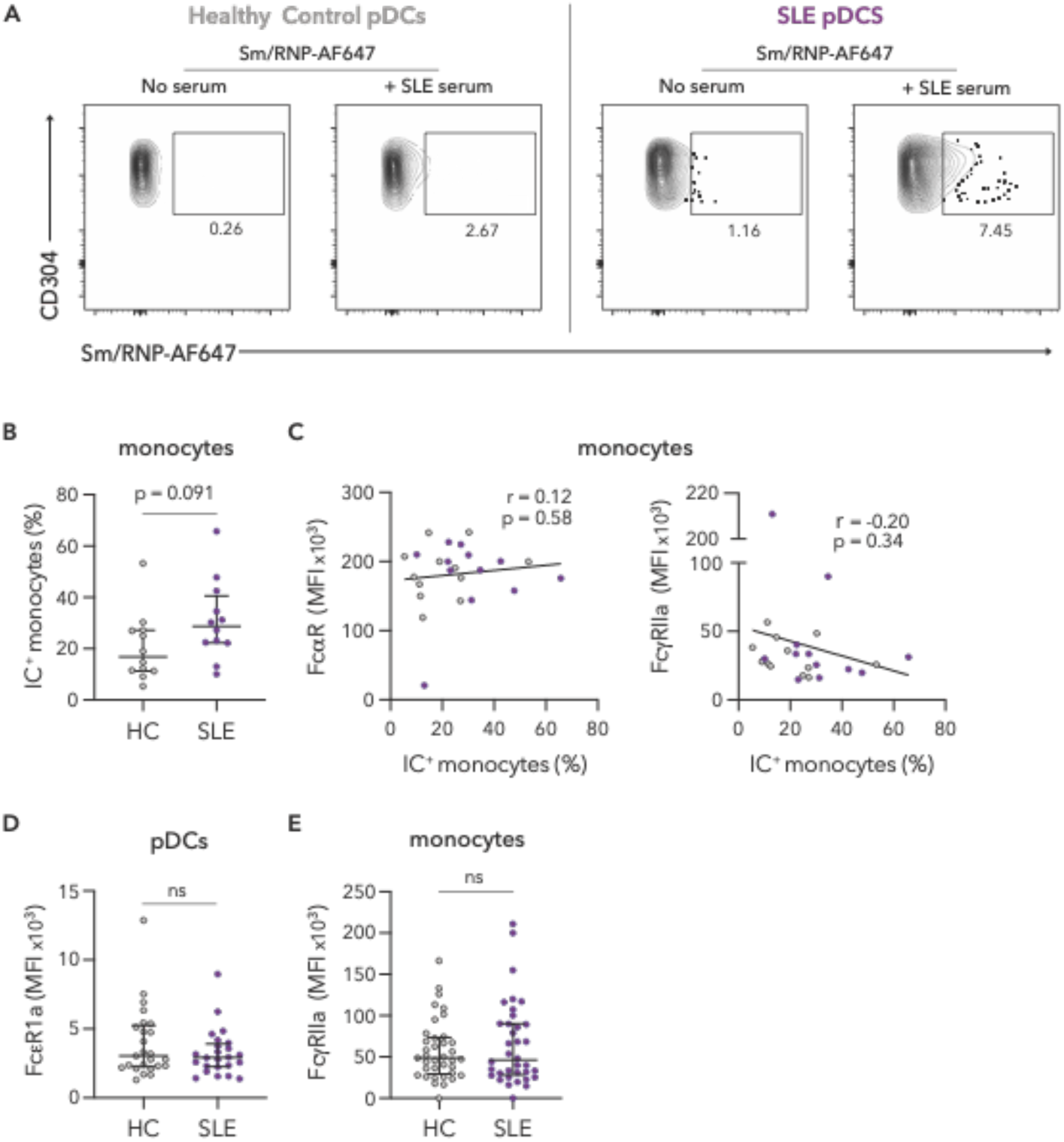
Supplementary figures relating to main Figure 7. **(A)** Representative flow plots for data shown in Fig. 7A in which thawed and rested PBMCs from individuals with SLE (n=12) and age-, race- and sex-matched healthy control (HC) subjects (n=12) were incubated for 3 hours with ICs generated from SLE serum and Sm/RNP-AF647. The first two flow plots show the percent of Sm/RNP-AF647+ healthy control pDCs when either Sm/RNP-AF647 is added alone (first) or in combination with SLE serum (second). The second two flow plots show the percent of Sm/RNP-AF647+ SLE pDCs when either Sm/RNP-AF647 is added alone (third) or in combination with SLE serum (fourth). **(B)** Thawed and rested PBMCs from individuals with SLE (n=12) and age-, race- and sex-matched healthy control subjects (n=12) were incubated for 3 hours with ICs generated from SLE serum and Sm/RNP-AF647. Percent of gated monocytes with ICs is shown. **(C)** PBMCs from the same subjects as in **(B)** were also stained for surface FcαR and FcγRIIa and correlations between IC^+^ monocytes and FcαR (*left*) or FcγRIIa (*right*) staining on monocytes are shown. Grey circles are pDCs from HC individuals and purple circles those from SLE individuals. **(D)** Cell surface expression of FcεRIa on gated pDCs from individuals with SLE (n=24) and age, race, and sex-matched HC subjects (n=24). **(E)** Cell surface expression of FcγRIIa on gated monocytes from individuals with SLE (n=36) and age, race, and sex-matched HC subjects (n=37). (**B, D** and **E**) Student’s t-tests **(C)** r = Pearson’s correlation coefficient.

**Supplementary Figure 8:**
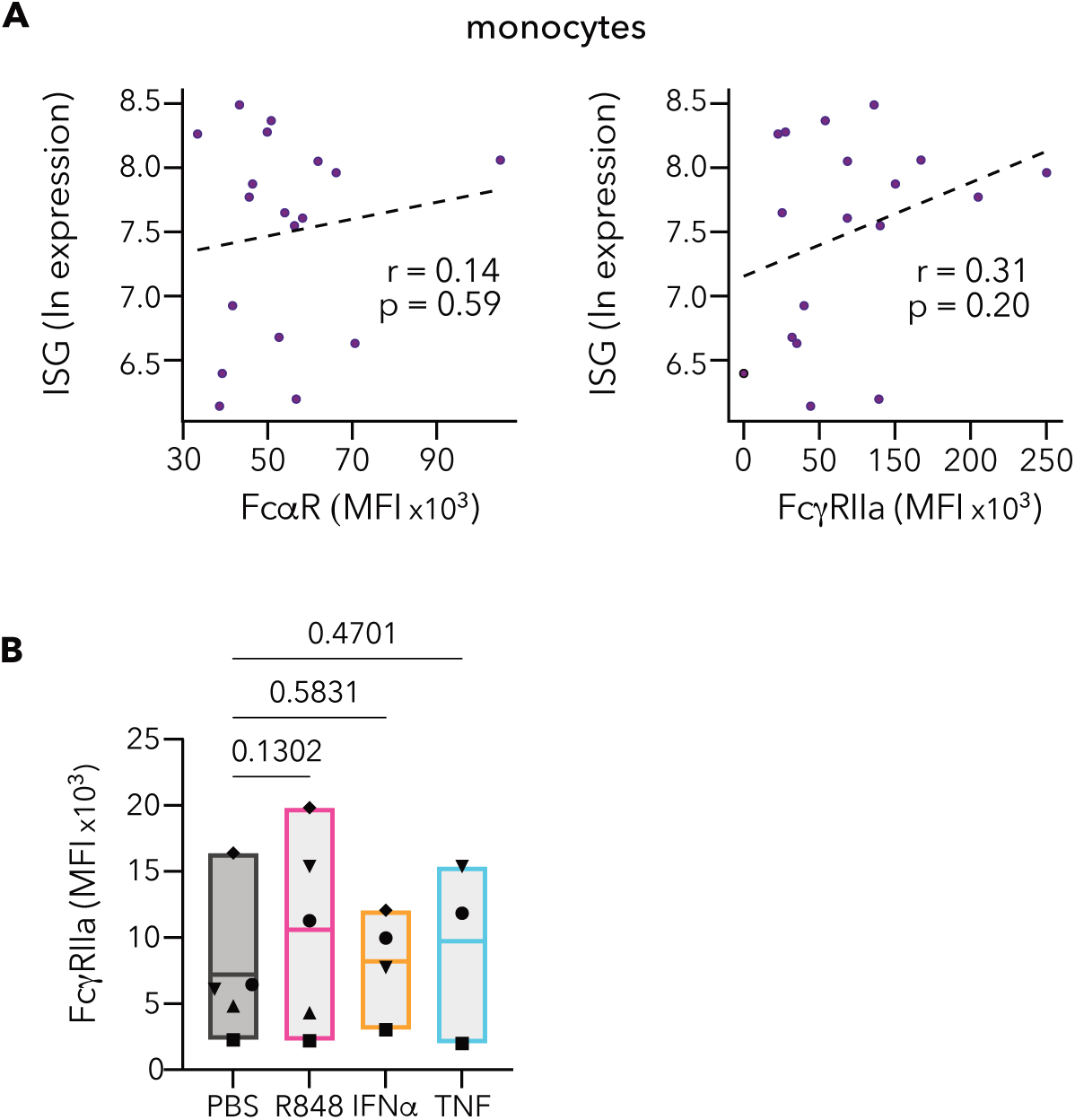
Supplementary figures relating to main Figure 8. **(A)** Correlation between cell surface expression of FcαR (*left*) and FcγRIIa (*right*) on gated monocytes and interferon-stimulated gene signature determined by RNA-seq performed on whole blood from 18 of the SLE subjects in Fig. 7C. **(B)** pDC FcγRIIa expression (geometric MFI) after 36 hr incubation with PBS control, TLR7 agonist R848, IFNα or TNF. (**A**) r = Pearson’s correlation coefficient. (**B**) One way ANOVA, repeated measures comparing all induction conditions to the PBS control.

**Supplementary Table 1:**
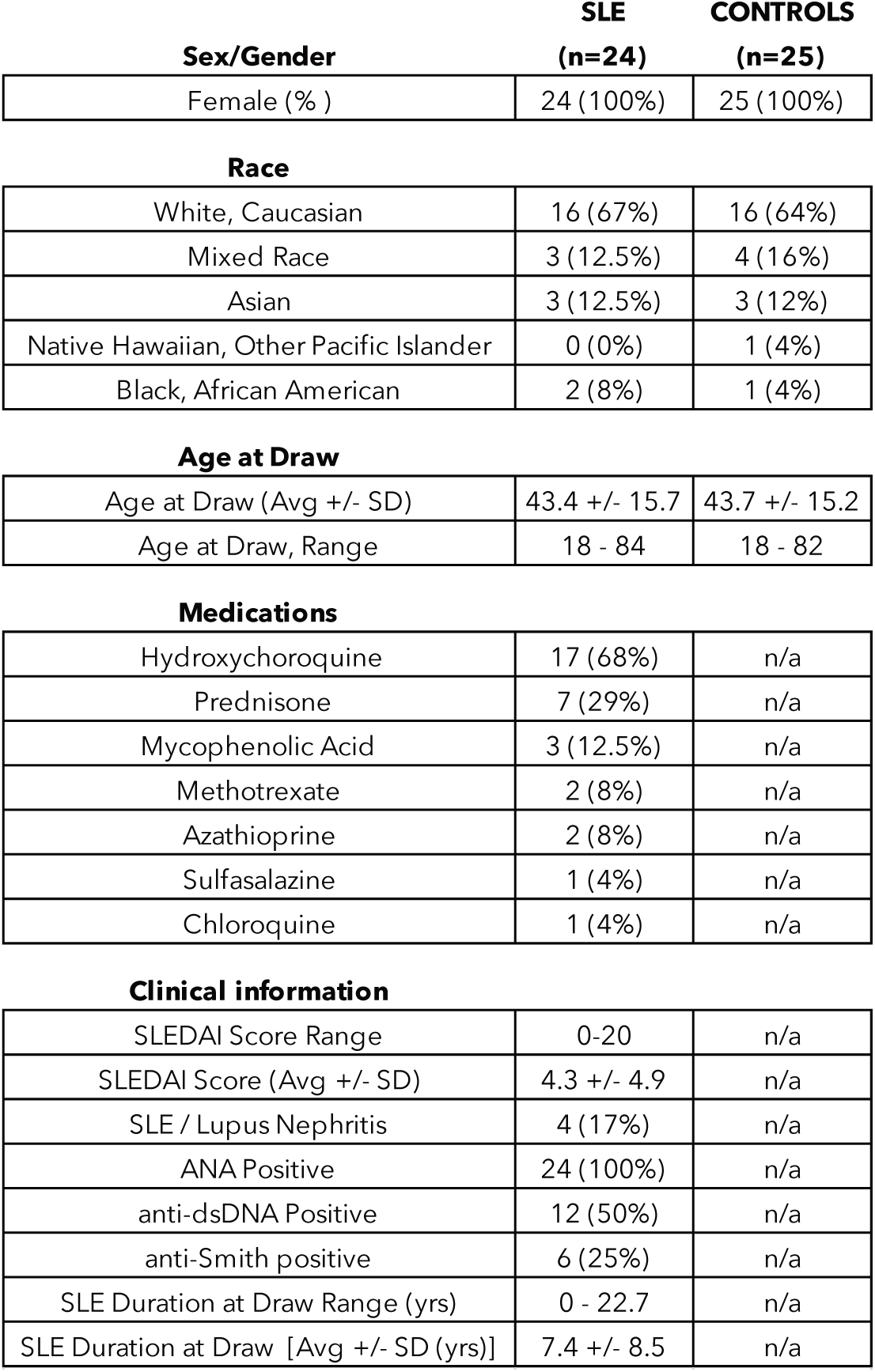
SLE and HC Participant Population Summary.

**Supplementary Table 2:**
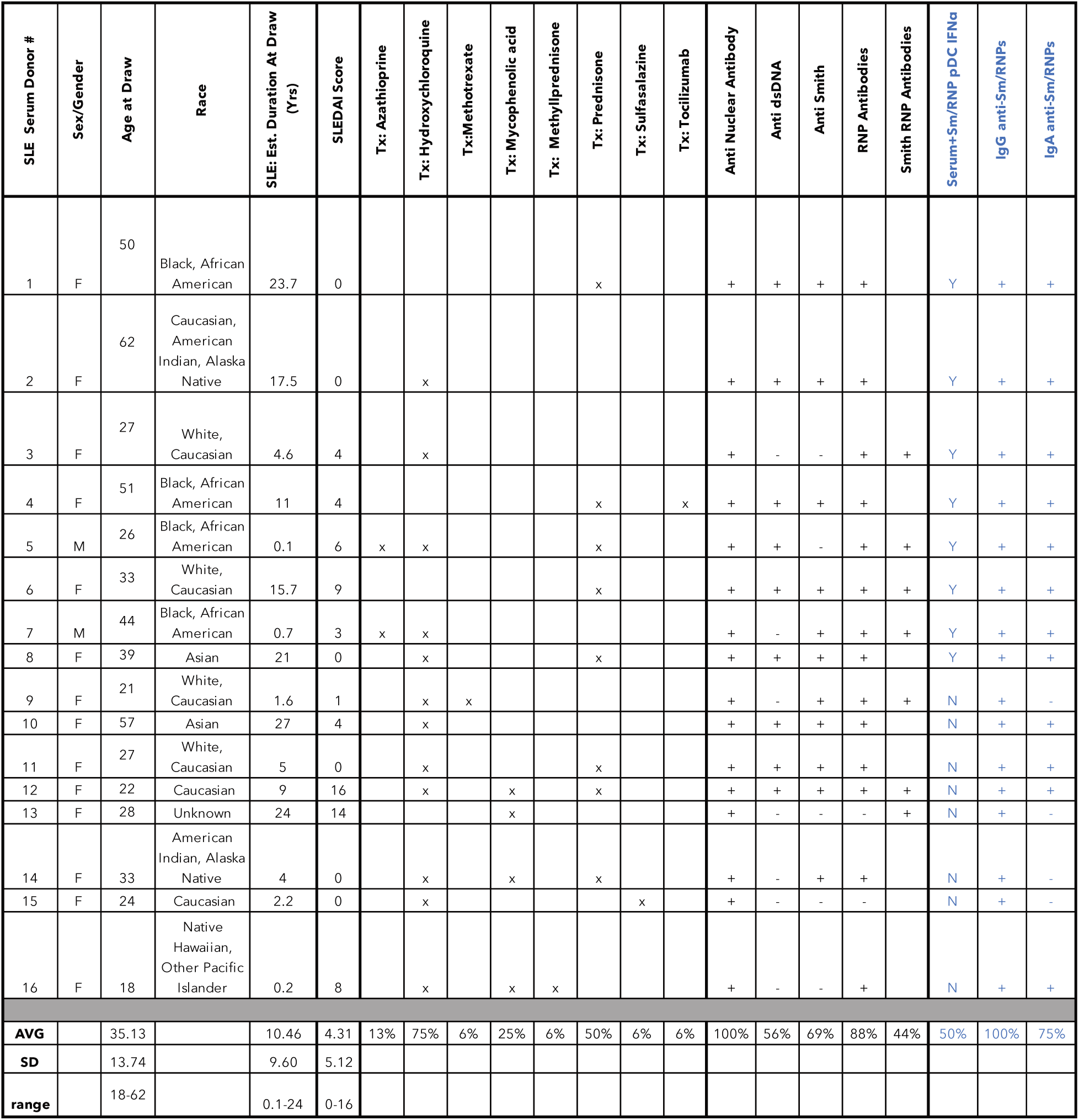
SLE Participant Population Detailed Summary.

**Supplementary Table 3:**
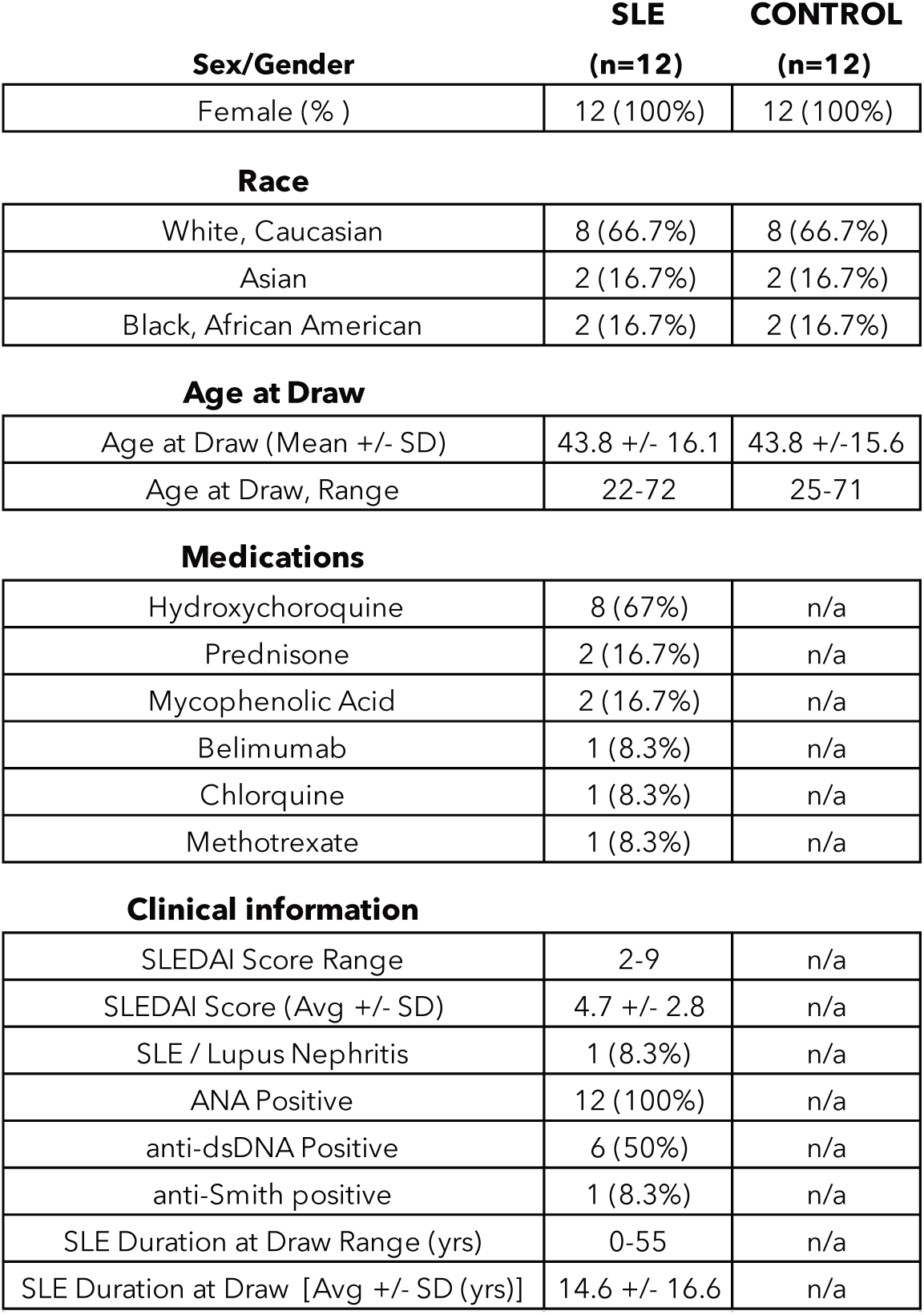
SLE and HC Participant Population Summary.

**Supplementary Table 4:**
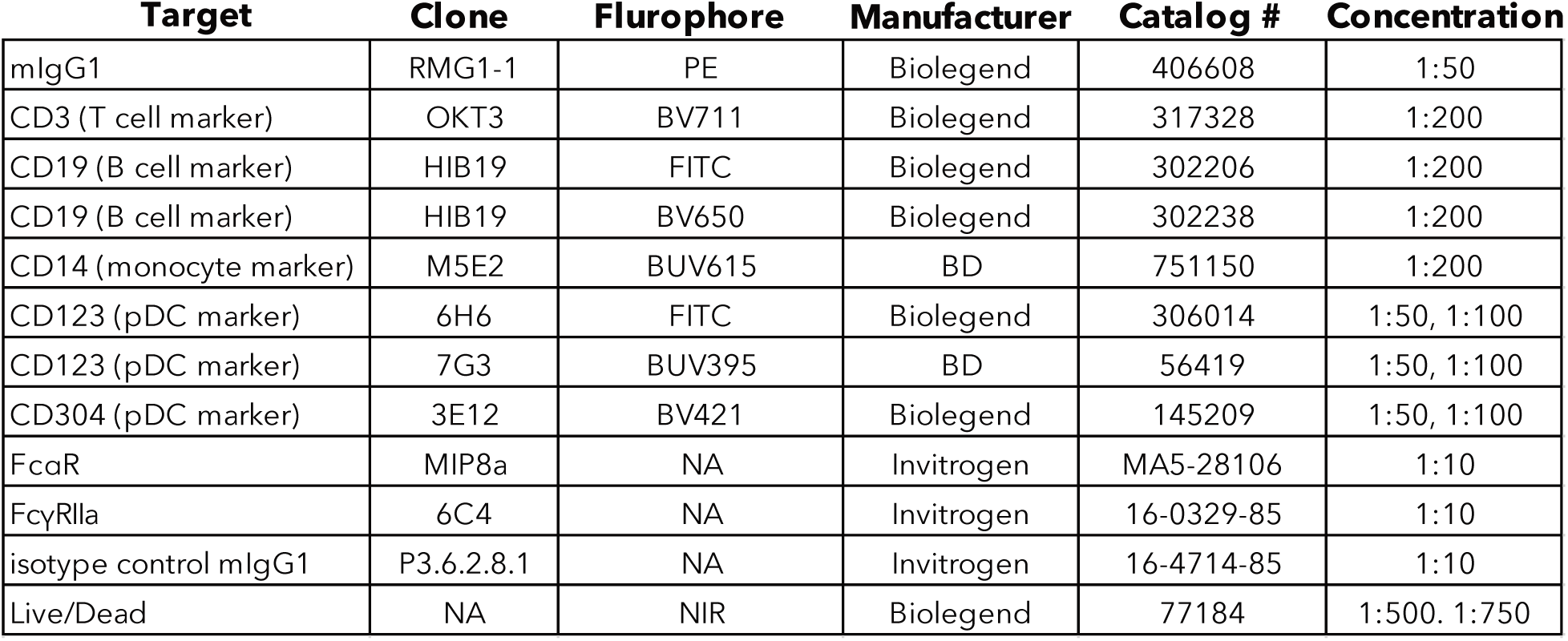
Flow Cytometry Antibody Summary.

