## Supplemental Figures and Tables for "Lupus IgA1 autoantibodies synergize with IgG to enhance pDC responses to RNA-containing immune complexes"

### Supplementary Figure 1

**A**

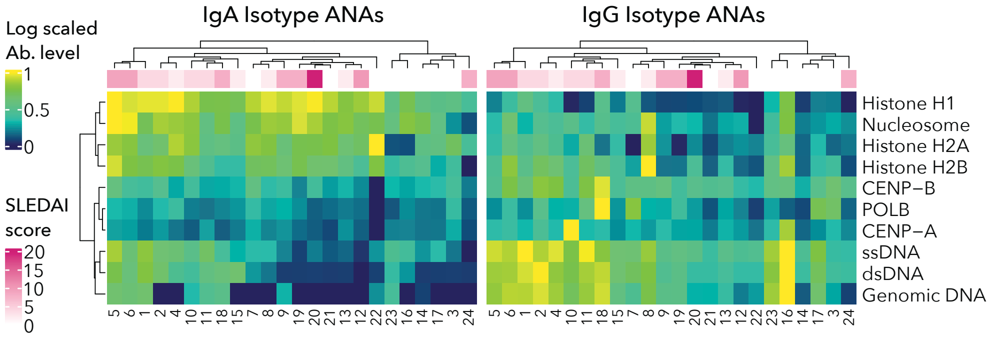

**B**

| Nuclear Antigen | Mean (NSI) IgA | Mean (NSI) IgG | Ratio (NSI) IgA/IgG | (r) ln IgA to IgG | (p) ln IgA to IgG |
|---|---|---|---|---|---|
| CENP-A | 48 | 915 | 0.05 | 0.13 | 0.5542 |
| CENP-B | 133 | 494 | 0.27 | 0.28 | 0.1781 |
| POLB | 154 | 836 | 0.18 | 0.39 | 0.0564 |
| **Genomic DNA** | **45** | **1111** | **0.04** | **0.51** | **0.0102** |
| Histone H1 | 2855 | 466 | 6.12 | 0.10 | 0.6458 |
| Histone H2A | 4017 | 1189 | 3.38 | -0.21 | 0.3351 |
| Histone H2B | 966 | 746 | 1.29 | 0.21 | 0.3356 |
| Nucleosome antigen | 3013 | 812 | 3.71 | 0.30 | 0.1608 |
| **dsDNA** | **272** | **3545** | **0.08** | **0.81** | **1.7E-06** |
| **ssDNA** | **1842** | **16628** | **0.11** | **0.88** | **1.4E-08** |
| **Jo-1** | **387** | **706** | **0.55** | **0.60** | **0.0018** |
| **La/SSB** | **217** | **643** | **0.34** | **0.66** | **0.0005** |
| **Ribo P. P0** | **462** | **1346** | **0.34** | **0.74** | **3.9E-05** |
| **Ribo P. P1** | **69** | **274** | **0.25** | **0.53** | **0.0082** |
| **Ribo P. P2** | **289** | **1658** | **0.17** | **0.71** | **3.9E-05** |
| **Ro/SSA (52 Kda)** | **1087** | **15321** | **0.07** | **0.66** | **0.0004** |
| Ro/SSA (60 Kda) | 110 | 458 | 0.24 | 0.40 | 0.0557 |
| Sm | 45 | 647 | 0.07 | 0.40 | 0.0545 |
| **Sm/RNP** | **139** | **2228** | **0.06** | **0.78** | **6.8E-06** |
| **SmD** | **68** | **672** | **0.10** | **0.52** | **0.0096** |
| **SmD1** | **143** | **1121** | **0.13** | **0.48** | **0.0167** |
| **U1-snRNP 68/70** | **300** | **5942** | **0.05** | **0.60** | **0.0018** |
| **U1-snRNP A** | **133** | **3272** | **0.04** | **0.54** | **0.0060** |
| **U1-snRNP B/B'** | **728** | **2431** | **0.30** | **0.52** | **0.0088** |
| **U1-snRNP C** | **846** | **3499** | **0.24** | **0.46** | **0.0224** |
| ssRNA | 67 | 299 | 0.22 | 0.25 | 0.2408 |

**Supplementary Figure 1: Supplementary figures relating to main Figure 1**

**(A)** Autoantibody profiling of serum from 24 SLE subjects. Heatmaps of log-scaled antibody binding for IgA (*left*) and IgG (*right*) isotype antibodies to 10 DNA-associated nuclear antigens. Each row corresponds to an antigen and each column corresponds to an individual subject. The numbering of subjects below heatmap correspond to the number in Fig. 1A. The color bar across the top indicates SLEDAI score for each donor at time of blood draw. Clustering of subjects and samples is based on similarity in scaled IgA levels, with the same clustering applied to the IgG heatmap. **(B)** Table shows mean normalized signal intensity (NSI) values for IgA and IgG isotype antibody, the ratio of IgA/IgG NSI values, as well the Pearson's correlation and p values for natural log transformed IgA and IgG NSI values.

#### Supplementary Figure 2

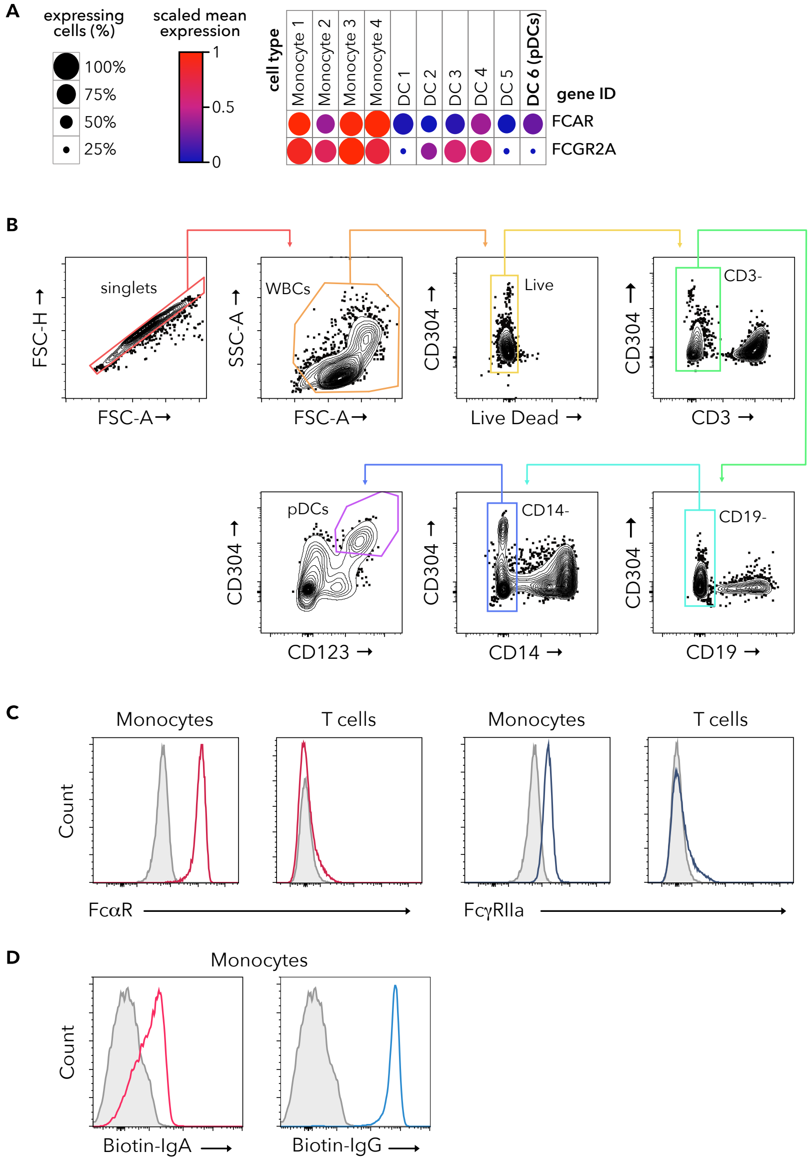

#### Supplementary Figure 2: Supplementary figures relating to main Figure 2

**(A)** *FCAR* and *FCGRIIA* mRNA expression in monocytes (1-4) and DC (1-6) subsets analyzed from publicly available single cell RNA sequencing data *(43)*. Color of the circles represent scaled mean expression levels and size of the circle corresponds to the percent of expressing cells for each group. **(B)** For all flow cytometry experiments gating for pDCs was done as shown. pDCs were identified as $CD3^-CD19^-CD14^-CD304^+CD123^+$ live single cells (*purple gate*). Monocyte and T cells were identified as $CD3^-CD19^-CD14^+$ and $CD19^-CD14^-CD3^+$ live single cells respectively. **(C)** PBMCs were analyzed by flow cytometry for surface FcαR (*red*) and FcγRIIa (*blue)* staining with isotype control staining in grey. The first two histograms show FcαR staining on gated monocytes (*first panel*) and T cells (*second panel*). The last two histograms show FcγRIIa on gated monocytes (*third panel*) and T cells (*fourth panel*) **(D)** Flow cytometry analysis showing aggregated biotinylated IgA (*red, left*) and IgG (*blue, right*) staining on monocytes as detected by labeled streptavidin. Gray histogram shows control streptavidin staining alone.

#### Supplementary Figure 3

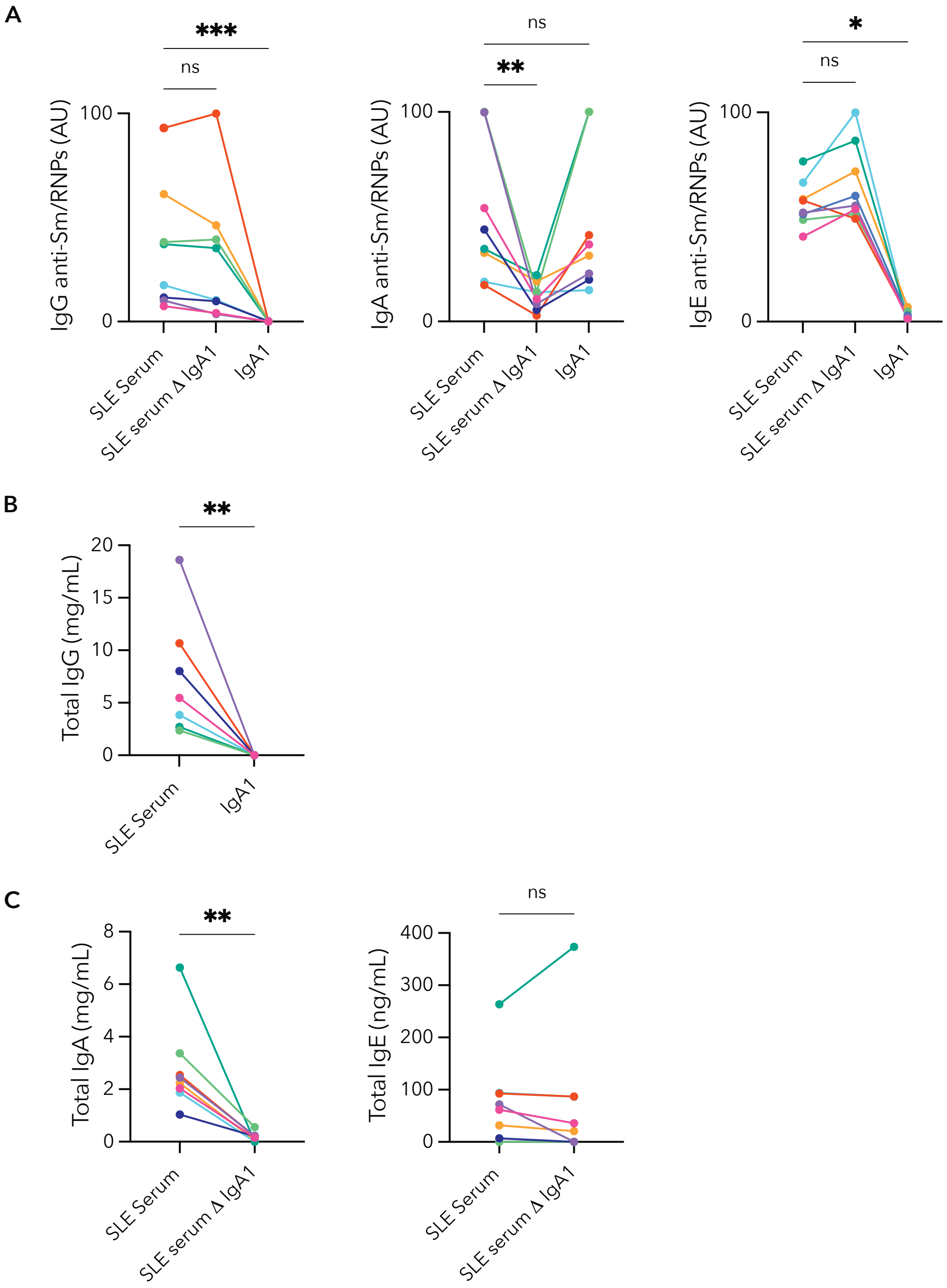

**Supplementary Figure 3: Supplementary figures relating to main Figure 3**

**(A)** IgG (*left*), IgA (*middle*) and IgE isotype (*right*) anti-Sm/RNP was measured in arbitrary units (AU) as described in methods. SLE serum, SLE serum ΔIgA1 and purified IgA1 were assayed for the 8 SLE serum donors used for all pDC IFNα production experiments. Each donor is color coded the same throughout this figure and in all other figures. **(B)** Total IgG in SLE serum and SLE IgA1. **(C)** Total IgA (*left*) in SLE serum and SLE serum ΔIgA1. Total IgE (*right*) in SLE serum and SLE serum ΔIgA1.

#### Supplementary Figure 4

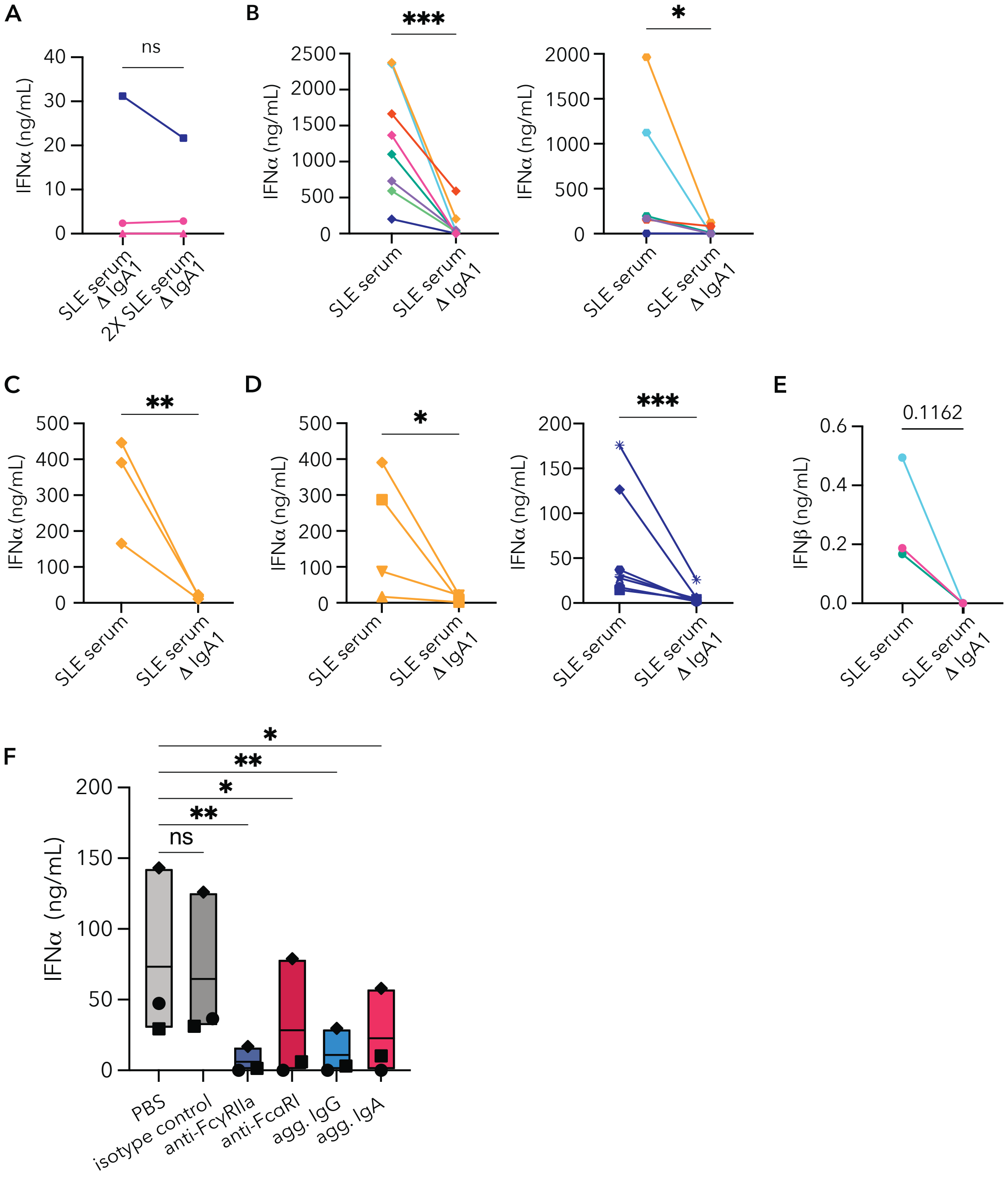

**Supplementary Figure 4: Supplementary figures relating to main Figure 4**

pDCs enriched from PBMCs freshly isolated from healthy control (HC) donor blood were incubated with immune complexes (ICs) generated with Sm/RNP RNA-containing nuclear antigens. **(A)** IFNα produced by pDCs after 24 hr incubation with Sm/RNP ICs generated with either SLE serum ΔIgA1 or twice the amount of SLE ΔIgA1 serum (2X SLE ΔIgA1 serum). **(B)** pDC IFNα production after 30-hour incubation with Sm/RNP ICs made with SLE serum or SLE serum ΔIgA1 from multiple SLE donors. Each line represents a distinct SLE serum donor, and each panel is from a different HC pDC donor distinct from the donor in Fig. 4C. **(C)** Paired analysis of pDC IFNα secretion with the same SLE serum donor and HC pDC donor assessed at 3 times over 7 months. **(D)** HC pDCs from multiple donors were stimulated with Sm/RNP

ICs made with SLE serum or SLE serum ΔIgA1 from one SLE serum donor (*left* same SLE serum donor as in Supplementary Fig. 4C, *right* same SLE donor from Fig. 4C). **(E)** Paired analysis of pDC IFNβ secretion after incubation with SLE serum or SLE serum ΔIgA1 ICs **(F)** Raw non-normalized data for experiment shown in Fig. 4F. HC pDCs were blocked with either PBS, isotype control antibody, monoclonal anti-FcγRIIa (6C4), monoclonal anti-FcαR (MIP8a), aggregated IgG or aggregated IgA for two hours before given Sm/RNP ICs generated with SLE serum. One way ANOVA with repeated measures ($p = 0.018$) with Bonferroni correction for multiple comparisons. Statistics for A, B and D-G are ratio paired t tests (* $p<0.05$, ** $p<0.01$, *** $p<0.001$.) SLE serum donors n=1 (C, D, F), n=2 (A), n=3 (E) n=6 (B *left*), n=8 (B *right*). HC pDC donors n=1 (B, C), n=3 (A, F), n=4 (D *left*), n=7 (D *right*).

#### Supplementary Figure 5

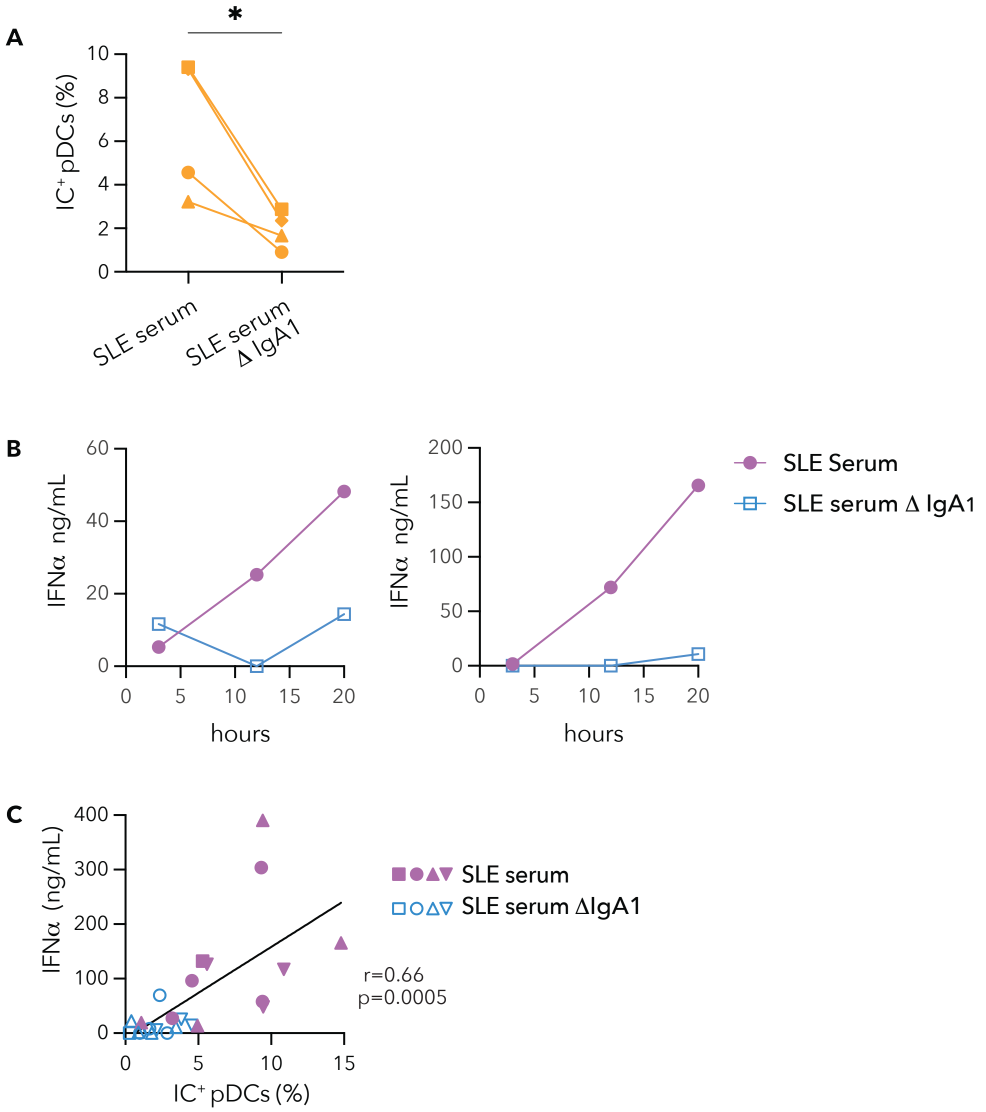

**Supplementary Figure 5: Supplementary figures relating to main Figure 5**

ICs were generated AlexaFluor647-labeled Sm/RNPs (Sm/RNP-AF647). **(A)** Paired analysis of percent $IC^+$ pDCs (n=4) after incubation with Sm/RNP-AF647 ICs generated with SLE serum or SLE serum ΔIgA1 from 1 SLE serum donor. **(B)** pDCs (n=1) were incubated with Sm/RNP-AF647 ICs generated with (n=2) SLE serum or SLE serum ΔIgA1 and IFNα secretion from replicate wells was measured at 3, 12 and 20 hours. Each graph shows data from a different SLE serum donor. **(C)** Correlation of 12 hour $IC^+$ pDCs (%) and 20 hour IFNα production for combined experiments in which pDCs were incubated with ICs generated with Sm/RNP-AF647 mixed with either SLE serum or SLE serum ΔIgA1. The combined data were generated with pDCs from n=4 HC pDC donors and n=4 SLE serum donors. r = Pearson's correlation coefficient.

#### Supplementary Figure 6

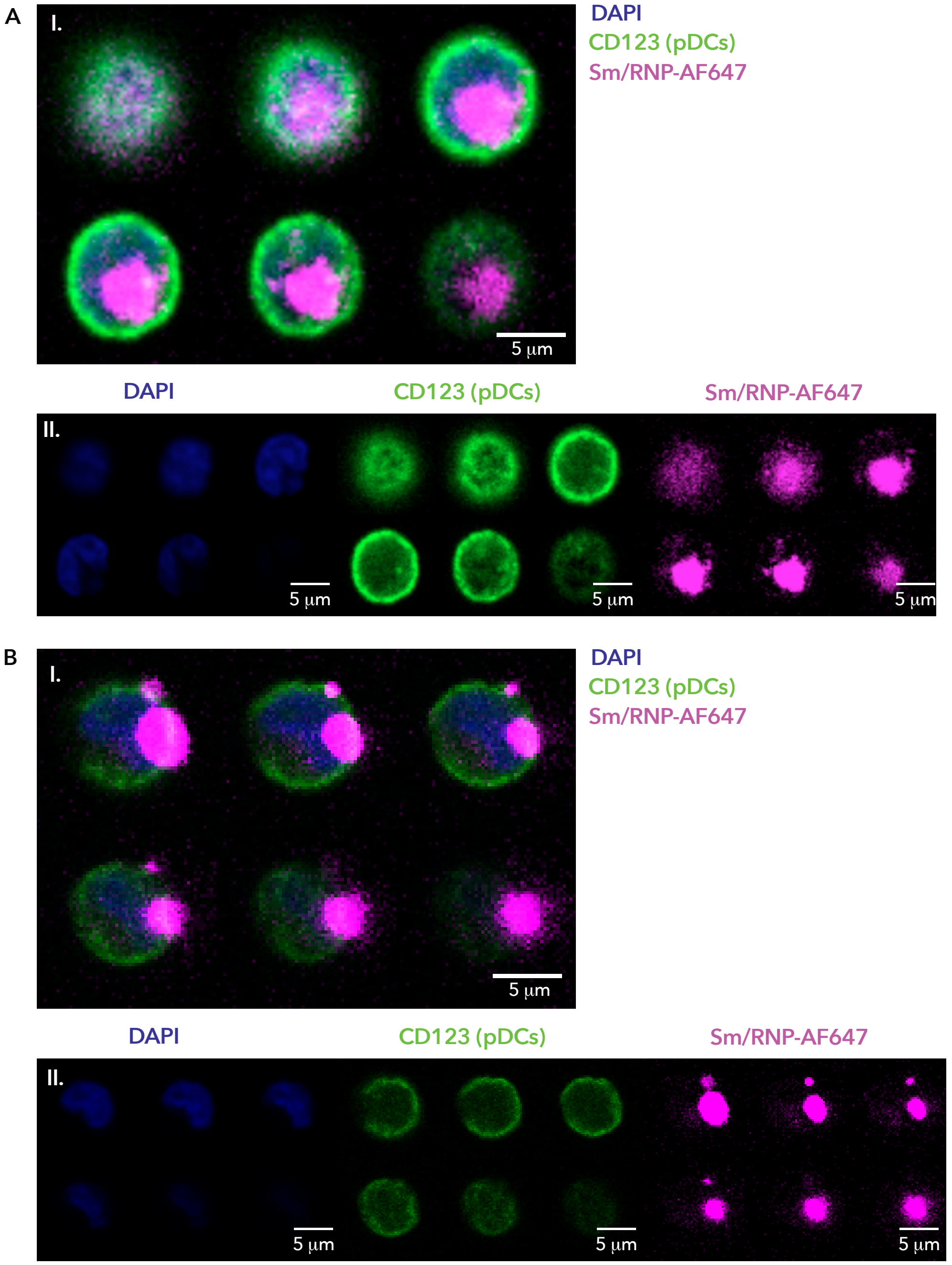

**Supplementary Figure 6: Representative images used to categorize Sm/RNP-AF647 SLE serum IC internalization and binding to pDCs**

Healthy control (HC) pDCs were incubated with ICs generated with Sm/RNP-AF647 alone or Sm/RNP-AF647 with SLE serum or SLE serum ΔIgA1 for 12 hours as in described in Fig. 6. **(A)** Representative example of a cell with large, internalized ICs. **(B)** Representative example of a cell with large ICs bound to the outside of the cell. For both **(A)** and **(B)** the first series of images (I.) shows the z-stack with all channels and second series of images (II.) show the z-stack for each individual channel; DAPI (*blue*), CD123 (*green*) and Sm/RNP-AF647 ICs (*magenta*).

**Supplementary Figure 7**

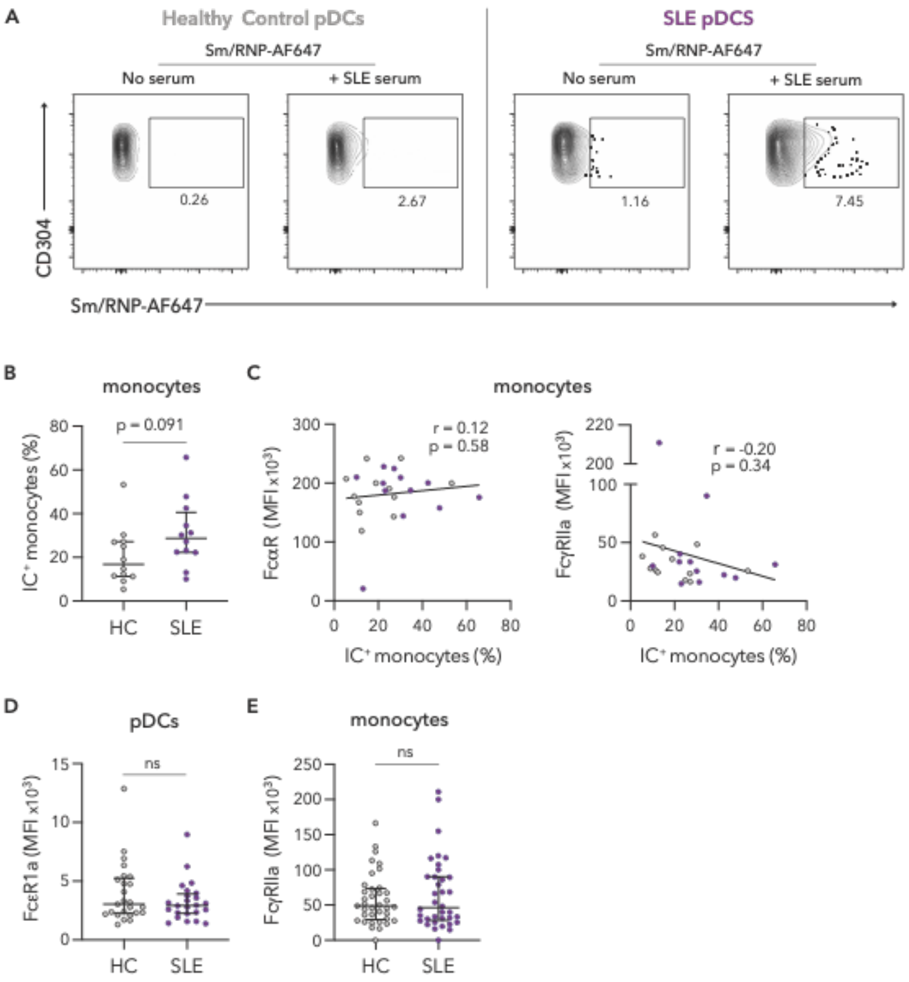

**Supplementary Figure 7: Supplementary figures relating to main Figure 7**

**(A)** Representative flow plots for data shown in Fig. 7A in which thawed and rested PBMCs from individuals with SLE (n=12) and age-, race- and sex-matched healthy control (HC) subjects (n=12) were incubated for 3 hours with ICs generated from SLE serum and Sm/RNP-AF647. The first two flow plots show the percent of Sm/RNP-AF647+ healthy control pDCs when either Sm/RNP-AF647 is added alone (first) or in combination with SLE serum (second). The second two flow plots show the percent of Sm/RNP-AF647+ SLE pDCs when either Sm/RNP-AF647 is added alone (third) or in combination with SLE serum (fourth). **(B)** Thawed and rested PBMCs from individuals with SLE (n=12) and age-, race- and sex-matched healthy control subjects (n=12) were incubated for 3 hours with ICs generated from SLE serum and Sm/RNP-AF647. Percent of gated monocytes with ICs is shown. **(C)** PBMCs from the same subjects as in **(B)** were also stained for surface FcαR and FcγRIIa and correlations between $IC^+$ monocytes and FcαR (*left*) or FcγRIIa (*right*) staining on monocytes are shown. Grey circles are pDCs from HC individuals and purple circles those from SLE individuals. **(D)** Cell surface expression of FcεRIa on gated pDCs from individuals with SLE (n=24) and age, race, and sex-matched HC subjects (n=24). **(E)** Cell surface expression of FcγRIIa on gated monocytes from individuals with SLE (n=36) and age, race, and sex-matched HC subjects (n=37). (**B, D** and **E**) Student's t-tests **(C)** r = Pearson's correlation coefficient.

**Supplementary Figure 8**

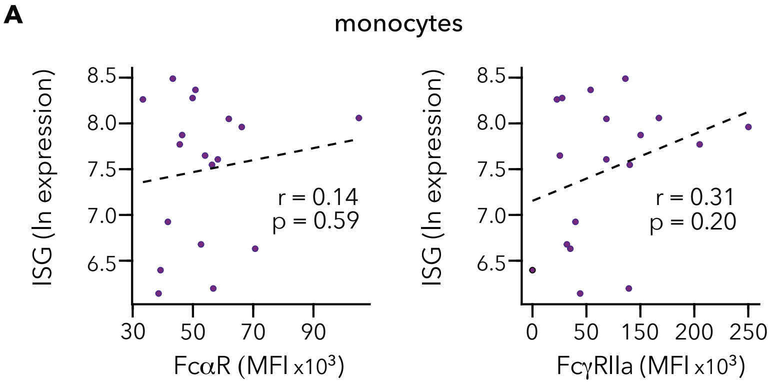

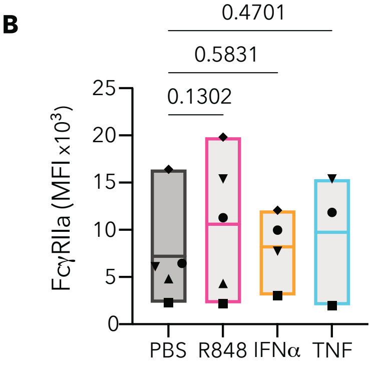

#### Supplementary Figure 8: Supplementary figures relating to main Figure 8

**(A)** Correlation between cell surface expression of FcαR (*left*) and FcγRIIa (*right*) on gated monocytes and interferon-stimulated gene signature determined by RNA-seq performed on whole blood from 18 of the SLE subjects in Fig. 7C. **(B)** pDC FcγRIIa expression (geometric MFI) after 36 hr incubation with PBS control, TLR7 agonist R848, IFNα or TNF. (**A**) r = Pearson's correlation coefficient. (**B**) One way ANOVA, repeated measures comparing all induction conditions to the PBS control.

#### Supplementary Table 1: SLE and HC Participant Population Summary

| Sex/Gender | SLE (n=24) | CONTROLS (n=25) |
|---|---|---|
| Female (% ) | 24 (100%) | 25 (100%) |

| Race | | |
|---|---|---|
| White, Caucasian | 16 (67%) | 16 (64%) |
| Mixed Race | 3 (12.5%) | 4 (16%) |
| Asian | 3 (12.5%) | 3 (12%) |
| Native Hawaiian, Other Pacific Islander | 0 (0%) | 1 (4%) |
| Black, African American | 2 (8%) | 1 (4%) |

| Age at Draw | | |
|---|---|---|
| Age at Draw (Avg +/- SD) | 43.4 +/- 15.7 | 43.7 +/- 15.2 |
| Age at Draw, Range | 18 - 84 | 18 - 82 |

| Medications | | |
|---|---|---|
| Hydroxychoroquine | 17 (68%) | n/a |
| Prednisone | 7 (29%) | n/a |
| Mycophenolic Acid | 3 (12.5%) | n/a |
| Methotrexate | 2 (8%) | n/a |
| Azathioprine | 2 (8%) | n/a |
| Sulfasalazine | 1 (4%) | n/a |
| Chloroquine | 1 (4%) | n/a |

| Clinical information | | |
|---|---|---|
| SLEDAI Score Range | 0-20 | n/a |
| SLEDAI Score (Avg +/- SD) | 4.3 +/- 4.9 | n/a |
| SLE / Lupus Nephritis | 4 (17%) | n/a |
| ANA Positive | 24 (100%) | n/a |
| anti-dsDNA Positive | 12 (50%) | n/a |
| anti-Smith positive | 6 (25%) | n/a |
| SLE Duration at Draw Range (yrs) | 0 - 22.7 | n/a |
| SLE Duration at Draw [Avg +/- SD (yrs)] | 7.4 +/- 8.5 | n/a |

### Supplementary Table 2: SLE Participant Population Detailed Summary

| SLE Serum Donor # | Sex/Gender | Age at Draw | Race | SLE: Est. Duration At Draw (Yrs) | SLEDAI Score | Tx: Azathioprine | Tx: Hydroxychloroquine | Tx:Methotrexate | Tx: Mycophenolic acid | Tx: Methyllprednisone | Tx: Prednisone | Tx: Sulfasalazine | Tx: Tocilizumab | Anti Nuclear Antibody | Anti dsDNA | Anti Smith | RNP Antibodies | Smith RNP Antibodies | Serum+Sm/RNP pDC IFNα | IgG anti-Sm/RNPs | IgA anti-Sm/RNPs |
|---|---|---|---|---|---|---|---|---|---|---|---|---|---|---|---|---|---|---|---|---|---|
| 1 | F | 50 | Black, African American | 23.7 | 0 | | | | | | x | | | + | + | + | + | | Y | + | + |
| 2 | F | 62 | Caucasian, American Indian, Alaska Native | 17.5 | 0 | | x | | | | | | | + | + | + | + | | Y | + | + |
| 3 | F | 27 | White, Caucasian | 4.6 | 4 | | x | | | | | | | + | - | - | + | + | Y | + | + |
| 4 | F | 51 | Black, African American | 11 | 4 | | | | | | x | | x | + | + | + | + | | Y | + | + |
| 5 | M | 26 | Black, African American | 0.1 | 6 | x | x | | | | x | | | + | + | - | + | + | Y | + | + |
| 6 | F | 33 | White, Caucasian | 15.7 | 9 | | | | | | x | | | + | + | + | + | + | Y | + | + |
| 7 | M | 44 | Black, African American | 0.7 | 3 | x | x | | | | | | | + | - | + | + | + | Y | + | + |
| 8 | F | 39 | Asian | 21 | 0 | | x | | | | x | | | + | + | + | + | | Y | + | + |
| 9 | F | 21 | White, Caucasian | 1.6 | 1 | | x | x | | | | | | + | - | + | + | + | N | + | - |
| 10 | F | 57 | Asian | 27 | 4 | | x | | | | | | | + | + | + | + | | N | + | + |
| 11 | F | 27 | White, Caucasian | 5 | 0 | | x | | | | x | | | + | + | + | + | | N | + | + |
| 12 | F | 22 | Caucasian | 9 | 16 | | x | | x | | x | | | + | + | + | + | + | N | + | + |
| 13 | F | 28 | Unknown | 24 | 14 | | | | x | | | | | + | - | - | - | + | N | + | - |
| 14 | F | 33 | American Indian, Alaska Native | 4 | 0 | | x | | x | | x | | | + | - | + | + | | N | + | - |
| 15 | F | 24 | Caucasian | 2.2 | 0 | | x | | | | | x | | + | - | - | - | | N | + | - |
| 16 | F | 18 | Native Hawaiian, Other Pacific Islander | 0.2 | 8 | | x | | x | x | | | | + | - | - | + | | N | + | + |
| **AVG** | | 35.13 | | 10.46 | 4.31 | 13% | 75% | 6% | 25% | 6% | 50% | 6% | 6% | 100% | 56% | 69% | 88% | 44% | 50% | 100% | 75% |
| **SD** | | 13.74 | | 9.60 | 5.12 | | | | | | | | | | | | | | | | |
| **range** | | 18-62 | | 0.1-24 | 0-16 | | | | | | | | | | | | | | | | |

### Supplementary Table 3: SLE and HC Participant Population Summary

| Sex/Gender | SLE (n=12) | CONTROL (n=12) |
|---|---|---|
| Female (% ) | 12 (100%) | 12 (100%) |
| **Race** | | |
| White, Caucasian | 8 (66.7%) | 8 (66.7%) |
| Asian | 2 (16.7%) | 2 (16.7%) |
| Black, African American | 2 (16.7%) | 2 (16.7%) |
| **Age at Draw** | | |
| Age at Draw (Mean +/- SD) | 43.8 +/- 16.1 | 43.8 +/-15.6 |
| Age at Draw, Range | 22-72 | 25-71 |
| **Medications** | | |
| Hydroxychoroquine | 8 (67%) | n/a |
| Prednisone | 2 (16.7%) | n/a |
| Mycophenolic Acid | 2 (16.7%) | n/a |
| Belimumab | 1 (8.3%) | n/a |
| Chlorquine | 1 (8.3%) | n/a |
| Methotrexate | 1 (8.3%) | n/a |
| **Clinical information** | | |
| SLEDAI Score Range | 2-9 | n/a |
| SLEDAI Score (Avg +/- SD) | 4.7 +/- 2.8 | n/a |
| SLE / Lupus Nephritis | 1 (8.3%) | n/a |
| ANA Positive | 12 (100%) | n/a |
| anti-dsDNA Positive | 6 (50%) | n/a |
| anti-Smith positive | 1 (8.3%) | n/a |
| SLE Duration at Draw Range (yrs) | 0-55 | n/a |
| SLE Duration at Draw [Avg +/- SD (yrs)] | 14.6 +/- 16.6 | n/a |

**Supplementary Table 4: Flow Cytometry Antibody Summary**

| Target | Clone | Flurophore | Manufacturer | Catalog # | Concentration |
|---|---|---|---|---|---|
| mIgG1 | RMG1-1 | PE | Biolegend | 406608 | 1:50 |
| CD3 (T cell marker) | OKT3 | BV711 | Biolegend | 317328 | 1:200 |
| CD19 (B cell marker) | HIB19 | FITC | Biolegend | 302206 | 1:200 |
| CD19 (B cell marker) | HIB19 | BV650 | Biolegend | 302238 | 1:200 |
| CD14 (monocyte marker) | M5E2 | BUV615 | BD | 751150 | 1:200 |
| CD123 (pDC marker) | 6H6 | FITC | Biolegend | 306014 | 1:50, 1:100 |
| CD123 (pDC marker) | 7G3 | BUV395 | BD | 56419 | 1:50, 1:100 |
| CD304 (pDC marker) | 3E12 | BV421 | Biolegend | 145209 | 1:50, 1:100 |
| FcαR | MIP8a | NA | Invitrogen | MA5-28106 | 1:10 |
| FcγRIIa | 6C4 | NA | Invitrogen | 16-0329-85 | 1:10 |
| isotype control mIgG1 | P3.6.2.8.1 | NA | Invitrogen | 16-4714-85 | 1:10 |
| Live/Dead | NA | NIR | Biolegend | 77184 | 1:500. 1:750 |
